# Cryo-EM reveals how Hsp90 and FKBP immunophilins co-regulate the Glucocorticoid Receptor

**DOI:** 10.1101/2023.01.10.523504

**Authors:** Chari M. Noddings, Jill L. Johnson, David A. Agard

**Affiliations:** Department of Biochemistry and Biophysics, University of California, San Francisco, San Francisco, CA 94143, USA; Department of Biological Sciences, University of Idaho, Moscow, ID 83844, USA

## Abstract

Hsp90 is an essential molecular chaperone responsible for the folding and activation of hundreds of ‘client’ proteins, including the glucocorticoid receptor (GR)^1-3^. Previously, we revealed that GR ligand binding activity is inhibited by Hsp70 and restored by Hsp90, aided by co-chaperones^4^. We then presented cryo-EM structures mechanistically detailing how Hsp70 and Hsp90 remodel the conformation of GR to regulate ligand binding^5,6^. *In vivo*, GR-chaperone complexes are found associated with numerous Hsp90 co-chaperones, but the most enigmatic have been the immunophilins FKBP51 and FKBP52, which further regulate the activity of GR and other steroid receptors^7-9^. A molecular understanding of how FKBP51 and FKBP52 integrate with the GR chaperone cycle to differentially regulate GR activation *in vivo* is lacking due to difficulties reconstituting these interactions. Here, we present a 3.01 Å cryo-EM structure of the GR:Hsp90:FKBP52 complex, revealing, for the first time, that FKBP52 directly binds to the folded, ligand-bound GR using three novel interfaces, each of which we demonstrate are critical for FKBP52-dependent potentiation of GR activity *in vivo*. In addition, we present a 3.23 Å cryo-EM structure of the GR:Hsp90:FKBP51 complex, which, surprisingly, largely mimics the GR:Hsp90:FKBP52 structure. In both structures, FKBP51 and FKBP52 directly engage the folded GR and unexpectedly facilitate release of p23 through an allosteric mechanism. We also reveal that FKBP52, but not FKBP51, potentiates GR ligand binding *in vitro*, in a manner dependent on FKBP52-specific interactions. Altogether, we reveal how FKBP51 and FKBP52 integrate into the GR chaperone cycle to advance GR to the next stage of maturation and how FKBP51 and FKBP52 compete for GR:Hsp90 binding, leading to functional antagonism.

## Introduction

Hsp90 is required for the functional maturation of 10% of the eukaryotic proteome^10^. Hsp90 ‘clients’ are enriched in signaling proteins and transcription factors, such as steroid hormone receptors (SHRs), making Hsp90 an important clinical target^1^. SHRs, which include GR, are hormone-regulated transcription factors that depend on Hsp90 for function throughout their lifetimes^7,11-15^. We previously established *in vitro* reconstitution of the ‘GR-chaperone cycle’, revealing that GR depends on Hsp90 for function due to constant inactivation of ligand binding by Hsp70 and subsequent reactivation by Hsp90^4^. In the GR-chaperone cycle, now understood in atomic detail through cryo-EM, GR ligand binding is regulated by a cycle of three distinct chaperone complexes^5,6^. In this chaperone cycle, GR is first inhibited by Hsp70 and Hsp40, then loaded onto Hsp90:Hop (Hsp70/Hsp90 organizing protein co-chaperone) forming an inactive ‘GR-loading complex’ (GR:Hsp70:Hsp90:Hop)^5^. Upon ATP hydrolysis by Hsp90, two Hsp70s and Hop are released, and p23 is incorporated to form an active ‘GR-maturation complex’ (GR:Hsp90:p23), restoring GR ligand binding with enhanced affinity^6^. The cryo-EM structures of the GR-loading complex and GR-maturation complex reveal Hsp70 and Hsp90 locally unfold and refold the GR LBD in a controlled manner to directly regulate ligand binding.

*In vivo*, additional Hsp90 co-chaperones are found associated with the GR-chaperone cycle, including the large immunophilins, FKBP51 and FKBP52^7^. FKBP51 and FKBP52 are peptidyl proline isomerases (PPIases) that contain an N-terminal FK1 domain with PPIase activity, an FK2 domain lacking PPIase activity, and a C-terminal TPR domain, which canonically binds the EEVD motifs at the C-termini of Hsp90 and Hsp70^16-19^. Additionally, the TPR domain contains a helical extension at the C-terminus (Helix 7e), which was previously described to bind the C-terminal domain (CTD) closed dimer interface of Hsp90^20,21^. Although FKBP51 and FKBP52 are 70% similar in sequence, these co-chaperones have antagonistic functional effects on GR *in vivo*^*8*^. FKBP51 inhibits GR ligand binding, nuclear translocation, and transcriptional activity, while FKBP52 potentiates each of these fundamental GR activities^22-32^. FKBP51 and FKBP52 have also been implicated in the regulation of all other SHRs^8,9^. Due to the critical importance of steroid hormone signaling in the cell, altered expression of FKBP51 and FKBP52 is associated with various endocrine-related disease states, including a wide variety of cancers, infertility, stress and anxiety disorders, and immune-related diseases^8,9,33^. Despite their importance, the absence of structures of FKBP co-chaperones bound to Hsp90:client complexes precludes a mechanistic understanding of how these co-chaperones integrate with Hsp90:client complexes to regulate client function or how to design selective small-molecule therapeutics^33-38^. Here we present a 3.01 Å cryo-EM structure of the GR:Hsp90:FKBP52 complex, revealing for the first time how FKBP52 integrates into the GR-chaperone cycle and directly binds to the active client, potentiating GR activity *in vitro* and *in vivo*. We also present a 3.23 Å cryo-EM structure of the GR:Hsp90:FKBP51 complex, revealing how FKBP51 competes with FKBP52 for GR:Hsp90 binding and demonstrating how FKBP51 can act a potent antagonist to FKBP52.

## Results

### GR:Hsp90:FKBP52 Structure Determination

The GR:Hsp90:FKBP52 complex was prepared by *in vitro* reconstitution of the complete GR-chaperone cycle. GR DBD-LBD (amino acids 418-777) (hereafter, GR for simplicity) with an N-terminal maltose-binding protein (MBP) tag was incubated with Hsp70, Hsp40, Hop, Hsp90, p23, and FKBP52, allowing GR to progress through the chaperone cycle to reach the GR:Hsp90:FKBP52 complex (Extended Data Fig. 1a,b). The complex was stabilized with sodium molybdate and then purified by affinity purification on MBP-GR followed by size exclusion chromatography and light crosslinking (Extended Data Extended Data Fig. 1a,b). A 3.01 Å cryo-EM reconstruction of the GR:Hsp90:FKBP52 complex was obtained using RELION and CryoSparc, with atomic models built using Rosetta (Fig. 1a,b; Extended Data Fig. 1e, 2). The structure revealed a fully closed, nucleotide bound Hsp90 dimer (Hsp90A and Hsp90B) complexed with a single GR and a single FKBP52, which occupied the same side of Hsp90 (Fig. 1a,b, ED. Fig. 3a). Despite using a multi-domain GR construct, only the GR LBD was visible in the density map.

**Figure 1.**
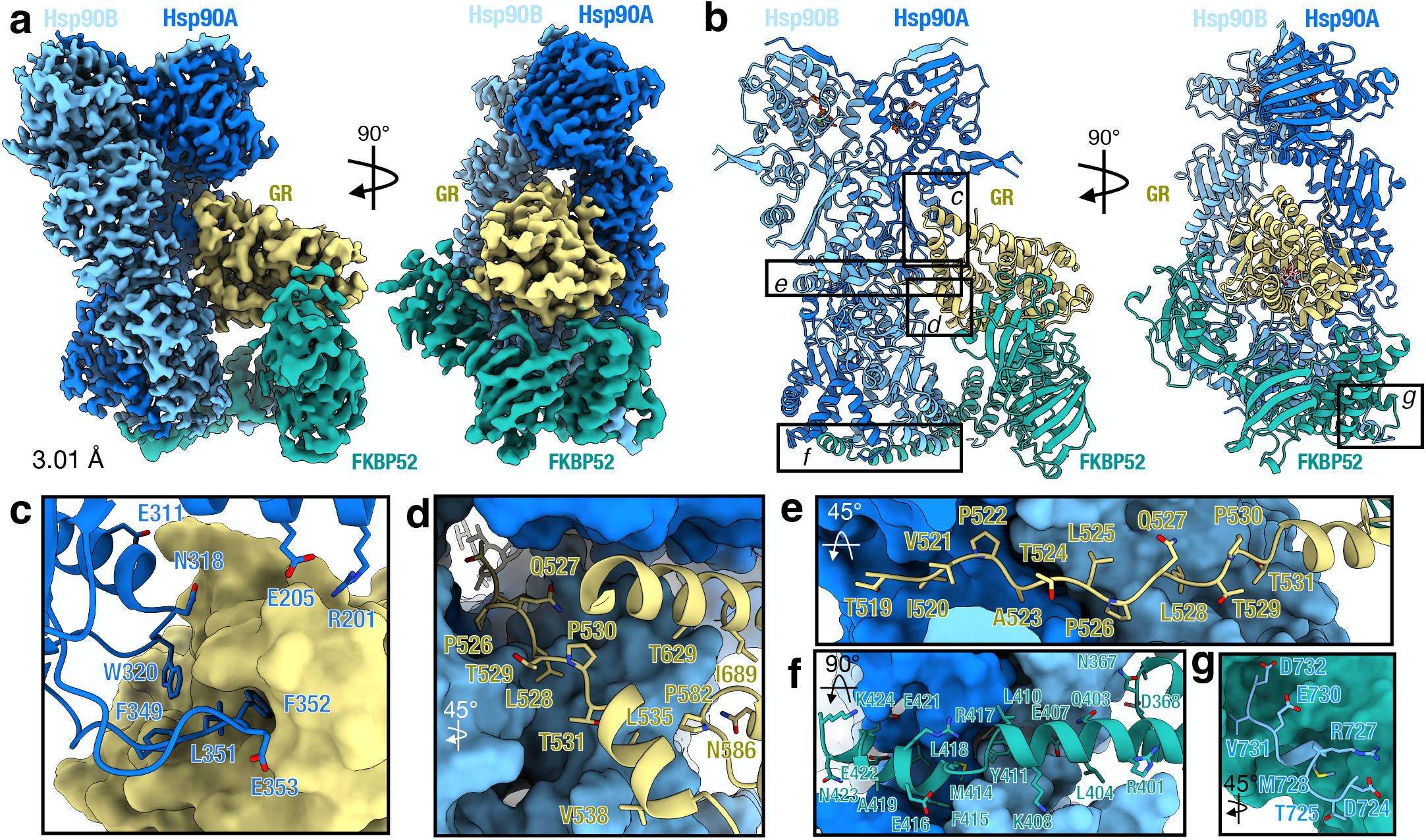
Architecture of the GR:Hsp90:FKBP52 complex. **a**, Composite cryo-EM map of the GR:Hsp90:FKBP52 complex. Hsp90A (dark blue), Hsp90B (light blue), GR (yellow), FKBP52 (teal). Color scheme is maintained throughout. **b**, Atomic model in cartoon representation with boxes corresponding to the interfaces shown in detail in **b-g. c**, Interface 1 of the Hsp90:GR interaction, depicting the Hsp90A Src loop (Hsp90A345-360) interacting with the GR hydrophobic patch. GR is in surface representation. **d**, Interface 2 of the Hsp90:GR interaction, depicting GR_Helix 1_ (GR532-539) packing against the entrance to the Hsp90 lumen. Hsp90A/B are in surface representation. **e**, Interface 3 of the Hsp90:GR interaction, depicting GR_pre-Helix 1_ (GR519-531) threading through the Hsp90 lumen. Hsp90A/B are in surface representation. **f**, Interface 1 of the Hsp90:FKBP52 interaction, depicting FKBP52 TPR H7e (FKBP52387-424) interacting with the Hsp90A/B CTD dimer interface. Hsp90A/B are in surface representation. **g**, Interface 2 of the Hsp90:FKBP52 interaction, depicting the Hsp90B MEEVD motif (Hsp90B700-706) binding in the helical bundle of the FKBP52 TPR domain. FKBP52 is in surface representation.

### Hsp90 Stabilizes the GR LBD in a Folded, Ligand-Bound State

In the GR:Hsp90:FKBP52 complex, GR adopts a fully folded, ligand-bound conformation (Extended Data Fig. 3b) distinct from that adopted in the GR:Hsp90:p23 maturation complex (discussed below). The folded GR is stabilized by Hsp90 at three major interfaces (Fig. 1c-e, Extended Data Fig. 3c-f): (1) Hsp90 Src loop:GR hydrophobic patch, (2) Hsp90 MD/CTD:GR Helix 1, and (3) Hsp90 lumen:GR pre-helix 1. In the first interface, the Hsp90A Src loop (Hsp90^345-360)^, flips out from the Hsp90 lumen, to interact with the previously described hydrophobic patch formed by GR Helices 9/10 and the GR C-terminus^6^ (approximately 767 Å^2^ of buried surface area (BSA)) (Fig. 1c, Extended Data Fig. 3c). Along the Src loop, Hsp90A ^F349,L351,F352,E353^ contact GR helices 9/10 and the conserved, solvent exposed Hsp90A^W320^ interacts with GR^F774^ on the GR C terminus. Notably, Hsp90A^W320,F349^ also make contact with GR in the GR-loading complex and GR-maturation complex, although at quite different locations^5,6^. Additionally, there are multiple hydrogen bonds formed between the Hsp90 NTD/MD to GR Helix 10 and the GR C-terminus (GR^K777^).

Interface 2 is comprised of Hsp90^Y604^ packing against GR Helix 1 (GR^532-539^) and Hsp90^Y627^ sticking into a hydrophobic pocket on GR formed by Helices 3, 4, and 9 (approximately 345 Å^2^ BSA) (Fig. 1d, Extended Data Fig. 3d,e). This GR hydrophobic pocket was previously identified in the androgen receptor (AR) as a druggable hydrophobic site (BF3)^39^. In interface 3, the unstructured GR pre-helix 1 region (GR^519-531^) is threaded through the closed Hsp90 lumen (approximately 758 Å^2^ BSA)(Fig. 1e, Extended Data Fig. 3f). Two hydrophobic residues on GR (GR^P522,P526^) occupy two hydrophobic pockets within the Hsp90 lumen. The interaction is further stabilized by multiple polar and hydrophobic interactions between GR pre-helix 1 and the Hsp90A/B amphipathic helical hairpin (Hsp90^606-628^) and Hsp90 MD.

### FKBP52 Interacts with the Closed Hsp90

FKBP52 engages the closed Hsp90 at three major interfaces (Fig. 1f,g, Extended Data Fig. 4a-c): (1) FKBP52 TPR H7e:Hsp90A/B CTDs, (2) FKBP52 TPR:Hsp90B MEEVD, and (3) FKBP52 TPR:Hsp90B CTD. In Interface 1, the extended TPR C-terminal H7e (FKBP52^387-424^) binds in a hydrophobic cleft formed by the Hsp90A/B CTDs at the closed dimer interface (approximately 1109 Å^2^ BSA)(Fig. 1f, Extended Data Fig. 4a). As compared to the crystal structure, H7e breaks at positions FKBP52^411-414^ to allow hydrophobic residues (FKBP52^L410,Y411,M414,F415,L418^) to flip into the hydrophobic cleft formed by the Hsp90 CTDs, consistent with the FKBP51 H7e:Hsp90 interaction observed by cryo-EM^20^. Mutating the corresponding conserved residues on FKBP51 H7e (FKBP51^M412,F413^ corresponding to FKBP52 ^M414,F415^) abolishes FKBP51:Hsp90 binding, indicating the importance of this binding site^20^. The interface is further stabilized by multiple hydrogen bonds and salt bridges from Hsp90A/B to H7e flanking the helix break (Extended Data Fig. 4a). Furthermore, a portion of the Hsp90B MEEVD linker (Hsp90B^700-706^) binds along FKBP52 H7e (Extended Data Fig. 4a).

In Interface 2, the C-terminal MEEVD peptide motif of Hsp90B binds in the FKBP52 TPR helical bundle (approximately 779 Å^2^ BSA) (Fig. 1g, Extended Data Fig. 4b), with multiple hydrogen bonds, salt bridges, and hydrophobic interactions, analogous to FKBP51:Hsp90^MEEVD^ structures^19,20^. However, the MEEVD peptide binds in an opposite orientation relative to the FKBP52:Hsp90^MEEVD^ crystal structure^18^, which may have been incorrectly modeled as previously suggested^19,40^. Interface 3 is comprised of the FKBP52 TPR helices 5/6 binding to the Hsp90B CTD, stabilized by multiple hydrogen bonds (approximately 193 Å^2^ BSA) (Extended Data Fig. 4c), also observed in the FKBP51:Hsp90 cryo-EM structure^20^. While the interactions between FKBP52 TPR/H7e:Hsp90 are conserved in the FKBP51:Hsp90 structure, the positions of the FKBP52 FK1 and FK2 domains are significantly altered (Extended Data Fig. 4d), owing to the presence of the bound GR client, as discussed below.

### FKBP52 Directly Interacts with GR, which is Functionally Important In Vivo

Unexpectedly, FKBP52 directly and extensively interacts with GR, with all three FKBP52 domains wrapping around GR, cradling the folded, ligand-bound receptor near the GR ligand-binding pocket (Fig. 2a). The tertiary structure within each FKBP52 domain closely matches isolated domains from FKBP52 crystal structures; however, the interdomain angles are significantly different (Extended Data Fig. 4d), likely owing to the extensive interaction with GR. There are three major interfaces between FKBP52 and GR (Fig. 2b-d): (1) FKBP52 FK1:GR, (2) FKBP52 FK2:GR, and (3) FKBP52 FK2-TPR linker:GR Helix 12.

**Figure 2.**
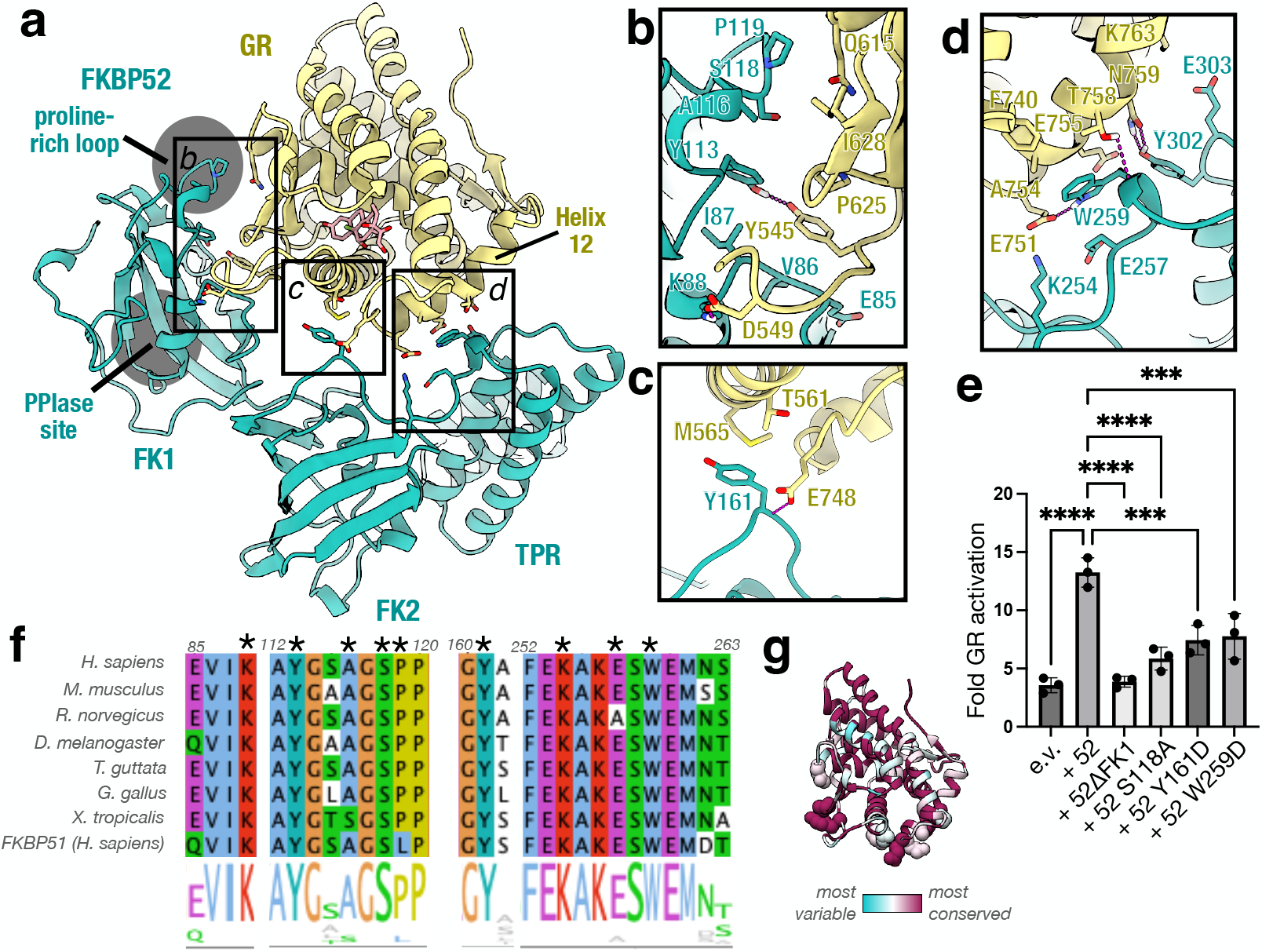
The GR:FKBP52 interaction and functional significance. **a**, Atomic model depicting the three interfaces between GR (yellow) and FKBP52 (teal) in the GR:Hsp90:FKBP52 complex. The FKBP52 proline-rich loop and PPIase site catalytic site are highlighted in gray. **b**, Interface 1 between GR (yellow) and the FKBP52 FK1 domain (teal), showing interacting side chains and hydrogen bonds (dashed pink lines). **c**, Interface 2 between GR (yellow) and the FKBP52 FK2 domain (teal), showing interacting side chains and hydrogen bonds (dashed pink lines). **d**, Interface 3 between GR (yellow) and the FKBP52 FK2-TPR linker (teal), showing interacting side chains and hydrogen bonds (dashed pink lines). **e**, GR activation assay in wild-type yeast strain JJ762 expressing FKBP52 (“52”) or FKBP52 mutants. The fold increase in GR activities compared to the empty vector (e.v.) control are shown (mean±SD). n=3 biologically independent samples per condition. Significance was evaluated using a one-way ANOVA (F_(6,14)_ = 67.82; p < 0.0001) with *post-hoc* Dunnett’s multiple comparisons test (n.s. P ≥ 0.05; * P ≤ 0.05; ** P ≤ 0.01; *** P ≤ 0.001). P-values: p(e.v. vs. 52) < 0.0001, p(52 vs. 52ΔFK1) < 0.0001, p(52 vs. 52 S118A) < 0.0001, p(52 vs. 52 Y161D) = 0.0001, p(52 vs. 52 W259D) = 0.0002. **f**, Sequence alignment of eukaryotic FKBP52 showing conserved residues involved in the GR:FKBP52 interaction (denoted by a black asterisk). The bottom aligned sequence is human FKBP51. The alignment is colored according to the ClustalW convention. **g**, GR protein sequence conservation mapped onto the GR atomic model from the GR:Hsp90:FKBP52 complex. Residue conservation is depicted from most variable (cyan) to most conserved residues (maroon).

In interface 1, FKBP52 FK1 interacts with a large surface on GR, canonically used for GR dimer formation, consisting of the GR post-helix 1 strand (helix 1-3 loop), helix 5, and β1,2 (approximately 280 Å^2^ BSA) (Fig. 2b). 3D variability analysis in CryoSparc reveals that the interaction between FKBP52 FK1 and GR is highly dynamic, even as the other FKBP52 domains (FK2, TPR) remain stably associated with GR (Supplemental Movies 1-2). At the FK1:GR interface, GR^Y545^ on the post-Helix 1 strand interacts with a hydrophobic surface formed by the FKBP52^81-88^ loop and forms a hydrogen bond with FKBP52^Y113^. Supporting this interaction, residues in the GR post-helix 1 strand (GR^544-546^) have previously been implicated in FKBP51/52-dependent regulation of GR activity^41,42^.

In addition, the FKBP52 proline-rich loop (β4-β5 loop or 80S loop) contacts GR Helix 5 and β1,2. 3D variability analysis in CryoSparc reveals that the proline-rich loop positioning is flexible, deviating from the position in the crystal structure (PDB ID: 4LAV)^43^ and adopting different interfaces with GR (Supplemental Movies 3-4). In the consensus 3D refinement map, the proline-rich loop adopts a position similar to the crystal structure, and FKBP52^A116,S118,P119^ interact with the tip of GR Helix 5 and β1,2. The FKBP52^P119L^ mutation has been shown to reduce GR and AR activation *in vivo*, while FKBP52^A116V^ has been shown to increase AR activation *in vivo*^29^. We also demonstrate that the FKBP52^S118A^ mutation significantly reduces FKBP52-dependent GR potentiation *in vivo* (Fig. 2e), further demonstrating the functional significance of this interaction site. In addition, S118 has been identified as a phosphorylation site on FKBP52, but not FKBP51 (qPTM database^44^) (possibly due to the unique adjacent proline on FKBP52 (P119) which could act as a signal for proline-directed protein kinases).

Phosphorylation at FKBP52 ^S118^ may help promote the interaction between the proline-rich loop and GR, which could also explain the large effect of the FKBP52^S118A^ mutation *in vivo*.

While the FKBP52 FK1 domain is known to have PPIase enzymatic activity, GR is not bound in the PPIase active site and accordingly, no GR prolines were found to have been isomerized compared to other GR structures (PDB ID: 1M2Z^45^, 7KRJ^6^). Consistent with this, mutation of GR prolines does not disrupt FKBP52-dependent regulation of GR^42^. Additionally, mutations that disrupt PPIase activity do not affect FKBP52-dependent GR potentiation *in vivo*^29^. Conversely, PPIase inhibitors have been shown to block the FKBP52-dependent potentiation of GR *in vivo*^*23*^. This can now be understood, as docking of PPIase inhibitors (FK506, rapamycin) into the PPIase active site demonstrate that the inhibitors would sterically block the FKBP52 FK1:GR interface (Extended Data Fig. 4e), which was previously hypothesized^23,29^.

Interface 2 is comprised of the FKBP52 FK2^Y161^ sticking into a shallow hydrophobic pocket formed by GR Helix 3 and the Helix 11-12 loop (GR^T561, M565,E748^) and a hydrogen bond between the FKBP52 backbone and GR^E748^ (approximately 125 Å^2^ BSA) (Fig. 2c). Supporting this interaction, we show that the FKBP52^Y161D^ mutation significantly reduces FKBP52-dependent GR potentiation *in vivo*, demonstrating the importance of this interaction (Fig. 2e). In interface 3, the solvent exposed, conserved W259 on the FKBP52 FK2-TPR linker makes electrostatic and hydrophobic interactions with GR Helix 12 (approximately 235 Å^2^ BSA) (Fig. 2d), which adopts the canonical agonist-bound position even in the absence of a stabilizing coactivator peptide interaction^45^ (Extended Data Fig. 3b). We show that the corresponding FKBP52^W259D^ mutation significantly reduces FKBP52-dependent GR potentiation *in vivo*, demonstrating the functional importance of this single residue (Fig. 2e). Interestingly, FKBP52^W259^ is also conserved in the FKBP-like co-chaperone XAP2 and a recent structure reveals XAP2 engages with an Hsp90-client using the analogous XAP2^W168^, suggesting this residue is critical more broadly for FKBP cochaperone:client engagement^46^. At interface 3, FKBP52^K254,E257,Y302,Y303^ make further polar interactions between the FK2-TPR linker and GR Helix 12 (Fig. 2d). While a significant portion of the GR Helix 12 co-activator binding site is available in the FKBP52-bound GR, the N-terminus of a co-activator peptide would sterically clash with the FKBP52 TPR based on the GR:co-activator peptide structure^45^ (Extended Data Fig. 5b). Thus, coactivator binding in the nucleus could help release GR from its complex with Hsp90:FKBP52. We also find that the residues at the FKBP52:GR interfaces are conserved across metazoans (Fig. 2f,g), in agreement with our results that single point mutations at each of the three FKBP52:GR interfaces has a significant effect on GR function *in vivo*.

### FKBP52 Advances GR to the Next Stage of Maturation

We previously described another GR-chaperone complex, the GR-maturation complex (GR:Hsp90:p23)^6^, which has important similarities and differences when compared to the GR:Hsp90:FKBP52 complex. Both the GR-maturation complex and the GR:Hsp90:FKBP52 complex are comprised of a closed Hsp90 dimer and a folded, ligand-bound GR (Fig. 3a). In the GR:Hsp90:FKBP52 complex, GR is rotated by approximately 45° relative to the GR-maturation complex (Fig. 3a). The Hsp90A Src loop interacts with the GR pre-Helix 1 strand in the maturation complex, but flips out to stabilize the rotated GR position in the GR:Hsp90:FKBP52 complex, by interacting with the GR hydrophobic patch (GR Helices 9/10)(Fig. 3b,c). In both complexes, the pre-Helix 1 strand of GR is threaded through the Hsp90 lumen; however, in the GR:Hsp90:FKBP52 complex, GR has translocated through the Hsp90 lumen by two residues, positioning two prolines (GR^P522,P526^) in the hydrophobic pockets in the Hsp90 lumen rather than two leucines (GR^L525,L528^) (Fig. 3d). This translocation positions the GR LBD further from Hsp90, likely allowing enough space for the observed GR rotation. The rotation of GR may also facilitate GR LBD dimerization, which is on pathway to activation (Extended Data Fig. 5c).

**Figure 3.**
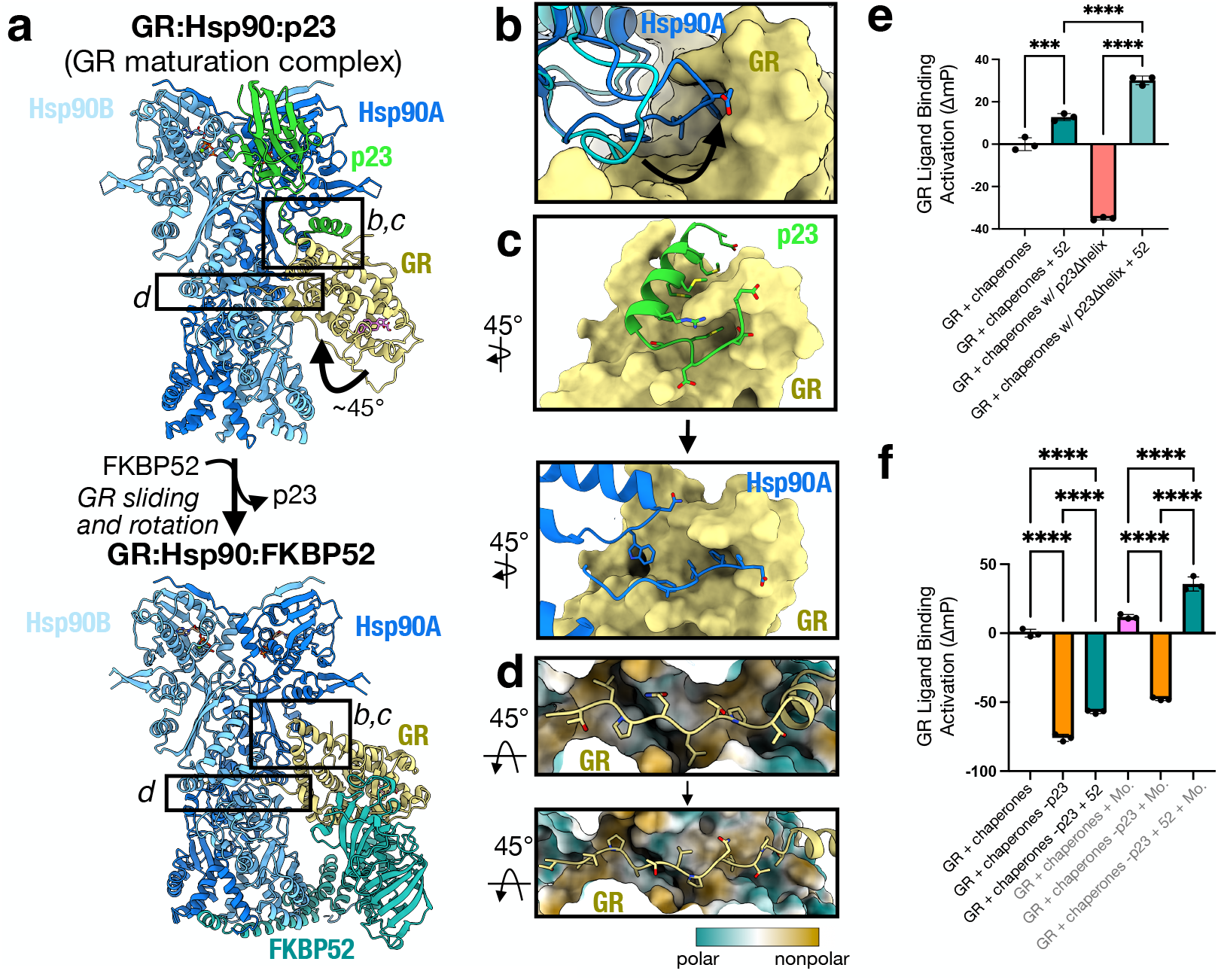
FKBP52 competes with p23 to bind GR:Hsp90. **a**, Atomic model of the GR-maturation complex (top) and the GR:Hsp90:FKBP52 complex (bottom) with boxes corresponding to the interfaces shown in detail in **b-d**. FKBP52 competes off p23 and re-positions GR at an approximately 45° rotated position. Hsp90A (dark blue), Hsp90B (light blue), GR (yellow), p23 (green), FKBP52 (teal). **b**, Position of the Hsp90A Src loop in the GR-maturation complex (Hsp90A, cyan) versus the GR:Hsp90:FKBP52 complex (Hsp90A, dark blue). The Hsp90A Src loop flips up in the GR:Hsp90:FKBP52 complex to interact with the hydrophobic patch on the rotated GR (yellow, surface representation). Hsp90A Src loop residues interacting with the GR hydrophobic patch are shown. **c**, Interface between the p23 tail-helix (green) and the GR hydrophobic patch (yellow, surface representation) in the GR-maturation complex (top). The p23 tail-helix is replaced by the Hsp90A Src loop (dark blue) in the GR:Hsp90:FKBP52 complex (bottom), which flip up to interact with the GR hydrophobic patch (yellow, surface representation). Interacting side chains are shown. **d**, Interaction between the GR_pre-Helix 1_ (GR_523-531_) threading through the Hsp90 lumen in the GR-maturation complex (top) versus the GR_pre-Helix 1_ (GR_519-531_) threading through the Hsp90 lumen in the GR:Hsp90:FKBP52 complex (bottom). Hsp90A/B are in surface representation colored by hydrophobicity. The GR_pre-Helix 1_ region translocates through the Hsp90 lumen by 2 residues in the transition from the GR-maturation complex to the GR:Hsp90:FKBP52 complex. Two hydrophobic residues on GR_pre-Helix 1_ (GR_L525,L528_ or GR_P522,P526_) remain bound in the Hsp90 lumen hydrophobic pockets in both complexes. **e**, Equilibrium binding of 10nM fluorescent dexamethasone to 100nM GR DBD-LBD with chaperones and FKBP52 (“52”). “Chaperones”= 15uM Hsp70, Hsp90, Hop, and p23 or p23Δhelix, 2uM Ydj1 and Bag-1. Significance was evaluated using a one-way ANOVA (F_(3,8)_ = 541.2; p < 0.0001) with *post-hoc* Šídák’s test (n.s. P ≥ 0.05; * P ≤ 0.05; ** P ≤ 0.01; *** P ≤ 0.001; **** P ≤ 0.0001). P-values: p(Chaperones vs. Chaperones) + 52 = 0.0002, p(Chaperones + 52 vs. Chaperones w/ p23Δhelix + 52) < 0.0001, p(Chaperones w/ p23Δhelix vs. Chaperones w/ p23Δhelix + 52) < 0.0001. **f**, Equilibrium binding of 10nM fluorescent dexamethasone to 100nM GR DBD-LBD with chaperones, FKBP52 (“52”), and sodium molybdate (“Mo”). “Chaperones”= 15uM Hsp70, Hsp90, Hop, and p23, 2uM Ydj1 and Bag-1. Significance was evaluated using a one-way ANOVA (F_(5,12)_ = 761.5; p < 0.0001) with *post-hoc* Šídák’s test (n.s. P ≥ 0.05; * P ≤ 0.05; ** P ≤ 0.01; *** P ≤ 0.001; **** P ≤ 0.0001). P-values < 0.0001 for each comparison.

Despite the translocation and rotation of GR, Hsp90 uses the same surfaces to bind GR (Hsp90B amphipathic helical hairpin, Hsp90A Src loop, Hsp90A^W320^); however, the GR contact surfaces are different.

### FKBP52 Competes with p23 for GR:Hsp90 Binding through Allostery

Surprisingly, FKBP52 competes with p23 to bind the GR:Hsp90 complex, although there is no direct steric conflict between FKBP52 and p23 binding (Fig. 3a). During 3D classification on the cryo-EM dataset, GR:Hsp90:p23 complexes were observed at low abundance; however, the GR:Hsp90:FKBP52 complexes showed no apparent p23 density (Extended Data Fig. 2), despite p23 being present at high concentration in the reconstitution mix. Furthermore, FKBP52 was found only associated with the rotated GR position, while the GR position in the p23-containing classes was only consistent with the GR-maturation complex. Thus, FKBP52 appears to specifically bind the rotated GR position, which is not compatible with p23 binding. This is consistent with mass spectrometry studies, demonstrating FKBP52 competes off p23 to form a stable GR:Hsp90:FKBP52 complex^47^. In the rotated GR position, the Hsp90A Src loop flips out of the Hsp90 lumen to bind the GR hydrophobic patch, which was previously engaged by the p23 tail-helix (Fig. 3a-c). Thus, rotation of GR dictates the accessibility of the hydrophobic patch to either Hsp90 or p23. FKBP52 stabilizes the rotated position of GR and therefore favors Hsp90 binding to GR over p23 and this in turn leads to p23 dissociation.

### FKBP52 Potentiates GR Ligand Binding In Vitro

To quantitatively assess the functional significance of FKBP52 on GR activation, we added FKBP52 to the *in vitro* reconstituted GR-chaperone cycle, using the GR DBD-LBD construct (residues 418-777) and monitored GR ligand binding, as previously described^4,6^.

Addition of FKBP52 to the GR chaperone cycle resulted in the enhancement of GR ligand binding above the already enhanced GR + chaperones control reaction at equilibrium (Fig. 3e), strongly suggesting FKBP52 potentiates the GR ligand binding affinity beyond the minimal chaperone mixture, consistent with reports *in vivo*^23^. We hypothesized that FKBP52 functions in a similar manner to the p23 tail-helix in stabilizing the ligand-bound GR. As previously described, removal of the p23 tail-helix (p23Δhelix) resulted in a decrease in GR ligand binding activity in the GR-chaperone system^6^; however, addition of FKBP52 to the reaction fully rescued GR ligand binding in the p23Δhelix background (Fig. 3e), suggesting FKBP52 functions in a similar manner to the p23 tail-helix in stabilizing the ligand-bound GR. Additionally, in the p23Δhelix background, FKBP52 potentiated ligand binding to a greater extent than in the wildtype p23 background. We hypothesize that removing the p23 tail-helix alleviates the competition between p23 and FKBP52, allowing p23 to remain bound to the GR:Hsp90:FKBP52 complex. Given that p23 is known to stabilize the closed Hsp90 conformation^6,48^, the enhanced ligand binding in the p23Δhelix background may be due to stabilization of closed Hsp90 by p23.

Interestingly, FKBP52 also affected GR ligand binding independent of Hsp90, with addition of FKBP52 to GR resulting in enhanced ligand binding, likely due to an Hsp90-independent chaperoning effect^16,49^ (Extended Data Fig. 5d).

### FKBP52 Functionally Replaces p23 In Vitro when Hsp90 Closure is Stabilized

Given that FKBP52 can functionally replace the p23 tail-helix, we wondered whether FKBP52 could also functionally replace p23 altogether. p23 is known to stabilize Hsp90 NTD closure through the globular p23 domain^6,48^ in addition to stabilizing the ligand-bound GR through the p23 tail-helix. Omitting p23 from the GR-chaperone cycle drastically reduces GR ligand binding, as previously described^4,6^. The addition of FKBP52 in place of p23 results in a modest increase in ligand binding but does not fully rescue ligand binding activity (Fig. 3f). We reasoned this could be due to the inability of FKBP52 to sufficiently stabilize Hsp90 closure, as previously suggested^50^. Therefore, we added molybdate to these reactions, which stabilizes NTD closure by acting as a γ-phosphate analogue in the Hsp90 NTD ATP-binding site^6,51^. Addition of molybdate to the reaction lacking p23 resulted in a small increase in GR ligand binding but did not fully rescue ligand binding activity. However, addition of molybdate to the reactions containing FKBP52 without p23 resulted in a full reactivation of ligand binding and even potentiated ligand binding over the control GR + chaperones reaction (Fig. 3f), much like with p23Δhelix. Thus, FKBP52 is able to functionally replace p23 if Hsp90 NTD closure is stabilized. Taken together, these results suggest FKBP52 can stabilize the ligand-bound GR, like p23, but cannot stabilize the closed Hsp90 NTD conformation, which requires p23.

### GR:Hsp90:FKBP51 Structure Determination

*In vivo* the interplay between FKBP52 and the highly similar FKBP51 have profound implications for GR activity. FKBP51 is functionally antagonistic to FKBP52-dependent potentiation of GR *in vivo*, thus the relative ratios of FKBP51 and FKBP52 dictate GR activity levels^23,28,52^. In order to understand mechanistically how FKBP51 antagonizes FKBP52, we prepared the analogous GR:Hsp90:FKBP51 complex (Extended Data Fig. 1c-e). We obtained a 3.23 Å cryo-EM reconstruction of GR:Hsp90:FKBP51 using RELION and CryoSparc, with atomic models built using Rosetta (Fig. 4a,b, Extended Data Fig. 6). Contrary to our expectations, the FKBP51-containing structure appears nearly identical to the FKBP52-containing structure. The GR:Hsp90:FKBP51 structure reveals a fully closed, nucleotide bound Hsp90 dimer complexed with a single GR and a single FKBP51, which occupy the same side of Hsp90 (Fig. 4a,b, Extended Data Fig. 7a). The FKBP51:Hsp90 interactions are analogous to the FKBP52:Hsp90 interactions, including the Hsp90B MEEVD:TPR interface and the Hsp90 CTD:TPR Helix 7e interface, also seen in the Hsp90:FKBP51:p23 structure^20^ (Fig. 4b, Extended Data Fig. 7b-d). The GR:Hsp90 interfaces are nearly identical when comparing the FKBP51 and FKBP52-containing complexes, including the Hsp90 Src loop:GR hydrophobic patch interface and the Hsp90 lumen:GR pre-Helix 1 strand interface (Fig. 4b, Extended Data Fig. 7e-g).

**Figure 4.**
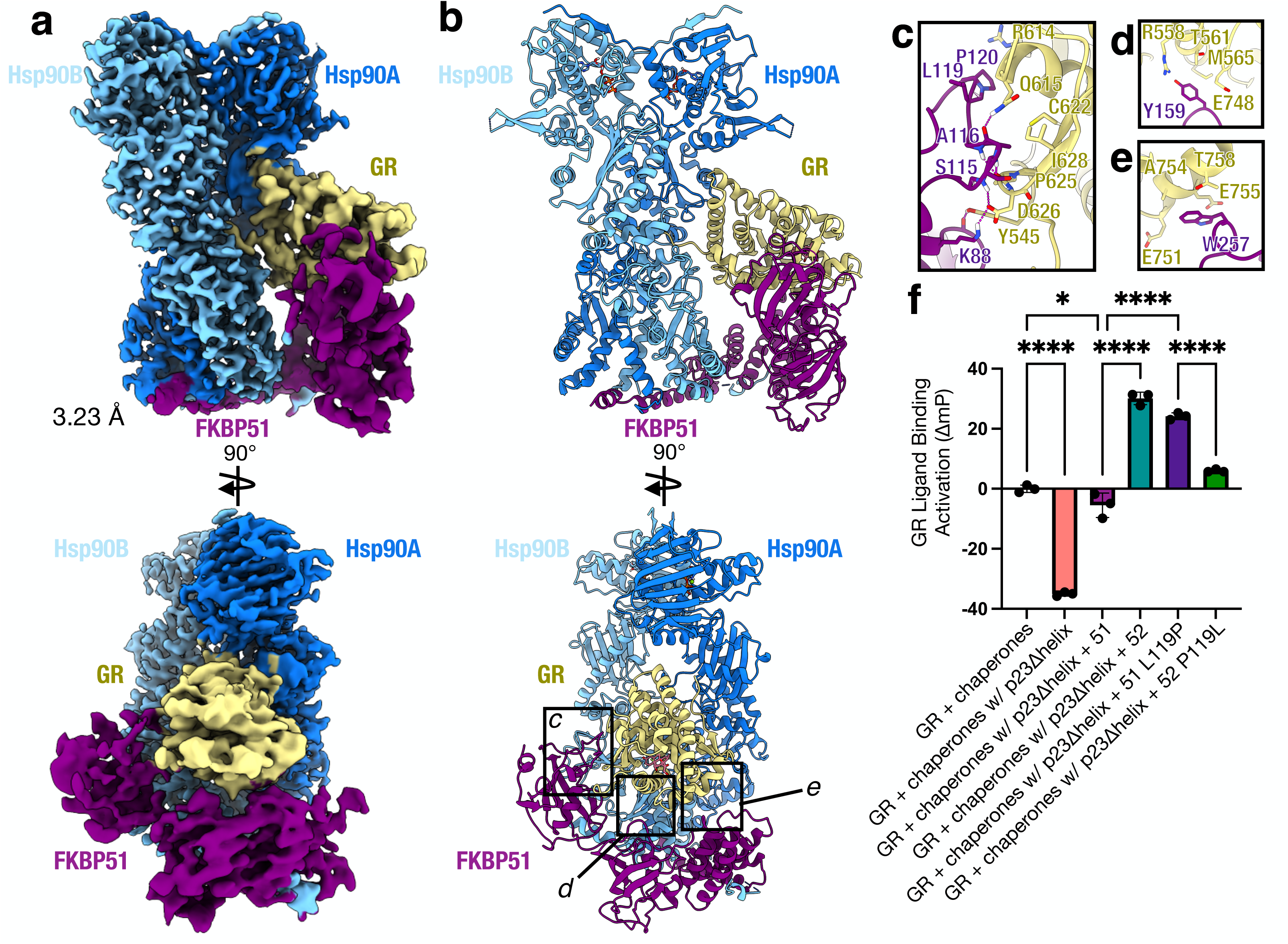
Architecture of the GR:Hsp90:FKBP51 complex. **a**, Composite cryo-EM map of the GR:Hsp90:FKBP51 complex. Hsp90A (dark blue), Hsp90B (light blue), GR (yellow), FKBP51 (purple). Color scheme is maintained throughout. **b**, Atomic model in cartoon representation with boxes corresponding to the interfaces shown in detail in **c-e. c**, Interface 1 between GR (yellow) and the FKBP51 FK1 domain (purple), showing interacting side chains and hydrogen bonds (dashed pink lines). **d**, Interface 2 between GR (yellow) and the FKBP51 FK2 domain (purple), showing interacting side chains and hydrogen bonds (dashed pink lines). **e**, Interface 3 between GR (yellow) and the FKBP51 FK2-TPR linker (yellow), showing interacting side chains and hydrogen bonds (dashed pink lines). **f**, Equilibrium binding of 10nM fluorescent dexamethasone to 100nM GR DBD-LBD with chaperones, FKBP51 (“51”), FKBP52 (“52”), or mutants. “Chaperones”= 15uM Hsp70, Hsp90, Hop, and p23 or p23Δhelix, 2uM Ydj1 and Bag-1. Significance was evaluated using a one-way ANOVA (F_(5,12)_ = 404.1; p < 0.0001) with *post-hoc* Šídák’s test (n.s. P ≥ 0.05; * P ≤ 0.05; ** P ≤ 0.01; *** P ≤ 0.001; **** P ≤ 0.0001). See **Methods** for p-values.

FKBP51 also directly binds GR in an analogous manner to FKBP52 (Fig. 4c-e). FKBP51 binds the folded, ligand-bound, rotated GR using the same three major interfaces (1) FKBP51 FK1:GR, (2) FKBP51 FK2:GR Helix 3, and (3) FKBP51 FK2-TPR linker:GR Helix 12. The GR:FKBP52 interaction residues are largely conserved for GR:FKBP51 (Fig. 2f). As with the FKBP52-containing structure, no GR prolines appear to be isomerized and the PPIase inhibitors rapamycin and FK506 sterically clash with the GR backbone. Interestingly, the small FKBP51-specific inhibitor, SAFit2 (PDB ID: 6TXX)^34,53^, does not clash with the GR backbone and may be accommodated in this complex with only side chain rotations, consistent with *in vivo* data from Baischew et al. 2022 (Extended Data Fig. 7h). Furthermore, the FKBP51 FK1 domain and FK1 proline-rich loop are highly dynamic, as revealed by CryoSparc 3D variability analysis, analogous to the FKBP52-containing structure (Supplemental Movies 5-6). However, in the GR:Hsp90:FKBP51 consensus map and corresponding atomic model, the FK1 domain contacts GR at a different angle relative to the GR:Hsp90:FKBP52 model; thus, the FK1:GR interface is distinct between the two complexes, specifically at the functionally important, but divergent, residue 119 in the proline-rich loop (FKBP51^L119^, FKBP52^P119^) (Fig. 2b, 3c)^29^, which we investigated further below.

### Functional Difference Between FKBP51 and FKBP52 Depends on Residue 119

To quantitatively assess the functional effect of FKBP51 on GR *in vitro*, we added FKBP51 to the GR-chaperone cycle and measured ligand binding activity. FKBP51 had no effect on the GR equilibrium value, unlike FKBP52, which potentiates GR ligand binding (Extended Data Fig. 8a). However, we found FKBP51 can functionally replace the p23 tail-helix or p23 (if molybdate is added), just as we observed with FKBP52 (Fig. 4f, Extended Data Fig. 8b).

However, FKBP51 does not potentiate GR ligand binding in any of these conditions, unlike FKBP52, recapitulating *in vivo* findings.

The residues responsible for the functional difference between FKBP51 and FKBP52 *in vivo* have been suggested to come from the proline-rich loop on the FK1 domain, specifically the divergent residue 119 (FKBP51^L119^, FKBP52^P119^)^29^. To assess whether this residue is responsible for the functional difference between FKBP51 and FKBP52 *in vitro*, we swapped residue 119 in FKBP51 and FKBP52 and added these mutants (FKBP51 L119P, FKBP52 P119L) to the *in vitro* reconstituted GR-chaperone cycle. We then measured ligand binding activity in the p23Δhelix background, where the largest potentiation due to FKBP52 is observed. Surprisingly, the residue 119 swapped mutants almost fully reversed the effects of FKBP51 and FKBP52 on GR— FKBP51 L119P potentiated GR ligand binding over the GR + chaperones control reaction, while FKBP52 P119L showed significantly less potentiation of ligand binding compared to wildtype FKBP52 (Fig. 4f). These results are consistent with the effects of the FKBP51/52 residue 119 swapped mutants *in vivo*^29^. Thus, residue 119 on the proline-rich loop provides a critical functional difference between the activities of FKBP51 and FKBP52 toward GR *in vitro* and *in vivo*, likely driven via the differential positioning of the loop seen in our consensus structures.

## Discussion

We present the first structures of the FKBP51 and FKBP52 co-chaperones bound to an Hsp90-client complex. The 3.01 Å GR:Hsp90:FKBP52 structure reveals that FKBP52 directly and extensively binds the client using three novel interfaces that stabilize the folded, ligand-bound conformation of GR. We show for the first time, that FKBP52 enhances GR ligand binding *in vitro*, consistent with *in vivo* reports, and that each of the three observed GR:FKBP52 interfaces is critical for FKBP52-dependent potentiation *in vivo*. We also provide a 3.23 Å GR:Hsp90:FKBP51 structure, unexpectedly demonstrating FKBP51 binds to the GR:Hsp90 complex in a similar manner to FKBP52. The FKBP51 interaction with closed Hsp90 is distinct from a previous NMR model^54^, but consistent with a recent cryo-EM structure demonstrating FKBP51 binds closed Hsp90^20^. Thus, these structures provide a molecular explanation for the functional antagonism between these two co-chaperones.

A recent study by Baischew et al. 2022 using *in vivo* chemical crosslinking validates our structures remarkably well, recapitulating all three major GR:FKBP51/52 contacts as well as the FKBP-mediated rotated GR position. Given that the *in vivo* crosslinking between GR and FKBP51/52 was performed in the absence of ligand, together our findings demonstrate that FKBP51 and FKBP52 bind the apo GR LBD in a similar, if not nearly identical manner to the ligand-bound GR LBD observed here in our structures. While our high-resolution reconstructions unambiguously contain ligand, apo GR:Hsp90:FKBP51/52 complexes likely exist in our dataset, but are less well-ordered (consistent with the results from the GR-maturation complex structure^6^). Baischew et al. 2022 provides further *in vivo* validation of our structural models by demonstrating FK506 inhibits FKBP51-dependent regulation of GR *in vivo*, while SAFit2 does not (Extended Data Fig. 4e, 7h), and that the FKBP51/52 FK1 domain is dynamically associated with GR relative to the other domains (Supplementary Movies 1-6).

Altogether, these studies complement each other extraordinarily well, demonstrating, for the first time, direct association of FKBP51 and FKBP52 with the GR LBD *in vivo* and *in vitro* at single-residue resolution.

Surprisingly, our structures also demonstrate that FKBP51 and FKBP52 compete with p23 to bind the GR:Hsp90 complex through an allosteric mechanism. Previous reports showed FKBP51 and p23 could simultaneously bind the closed Hsp90 in the absence of client^20^. We demonstrate that the position of the client can dictate which co-chaperone is bound, with the FKBPs and p23 binding to distinct GR positions. FKBP51 and FKBP52 both stabilize a rotated position of GR relative to the GR-maturation complex. One functional consequence of this rotated position may be to promote GR dimerization, which is a required step in GR activation. The rotated GR position relieves this steric hindrance to dimerization in the GR-maturation complex and would allow the GR LBD to dimerize once FKBP51/52 release (Extended Data Fig. 5c). Indeed, a previous report has suggested FKBP51/52 promote AR dimerization *in* vivo^55^, raising the possibility that the FKBPs promote this next step in SHR maturation.

Our structures also contribute to an emerging theme in which Hsp90 cochaperones bind to distinct Hsp90 conformations, while simultaneously binding to specific client conformations^5,6,46,51,56^. FKBP51 and FKBP52 each wrap around the folded, ligand-bound client using all three FKBP domains, while the FKBP TPR Helix 7e binds the closed Hsp90 conformation. The Helix 7e is found in many TPR-containing co-chaperones^20^, however, our structures, along with others, reveal the TPR Helix 7e can bind Hsp90 in distinct positions due to sequence divergence of H7e at the Hsp90 binding site^46,56^. Although the FKBPs directly contact GR, they do not appear to isomerize GR prolines or engage the GR NLS1 (nuclear localization signal 1) (GR^467-505^)^57^ to regulate GR activity, as previously hypothesized^13,58-60^

While FKBP51 binds similarly to FKBP52, competing with p23 and stabilizing the rotated GR, we find that FKBP51 does not significantly enhance GR ligand binding *in vitro*, like FKBP52, consistent with *in vivo* reports^23 27^. Interestingly, we find residue 119 on FKBP51/52 is critical for enhancement of ligand binding *in vitro*, also consistent with *in vivo* reports^29^. NMR studies have found the proline at residue 119 on FKBP52 decreases dynamics of the proline-rich loop (also called 80S loop or β4-β5 loop) relative to the leucine at FKBP51^61^. Analysis of dynamics of our structures using 3D variability analysis demonstrates that the proline-rich loop is highly dynamic in its interaction with GR. Thus, the dynamics of this loop may dictate the specificity and/or stability of this interaction, leading to distinct regulation of GR activity.

Based on our structures of the GR:Hsp90:FKBP51 and GR:Hsp90:FKBP52 complexes, we propose additional steps in the GR-chaperone cycle that account for FKBP51/52 incorporation and subsequent regulation of GR activity in the cell (Fig. 5). In the cytosol, GR cycles between Hsp70 and Hsp90, which locally unfold and refold GR to directly control ligand binding, as previously described^4-6^. Once the folded, ligand-bound GR reaches the GR-maturation complex (GR:Hsp90:p23), either FKBP51 or FKBP52 can bind the complex and compete with p23 to advance GR to the next stage of maturation. Given that the folded GR is strongly stabilized and tightly associated with Hsp90 and the FKBPs, we suggest that it is unlikely that ligand binding or unbinding happens in the context of FKBP-bound complexes.

**Figure 5.**
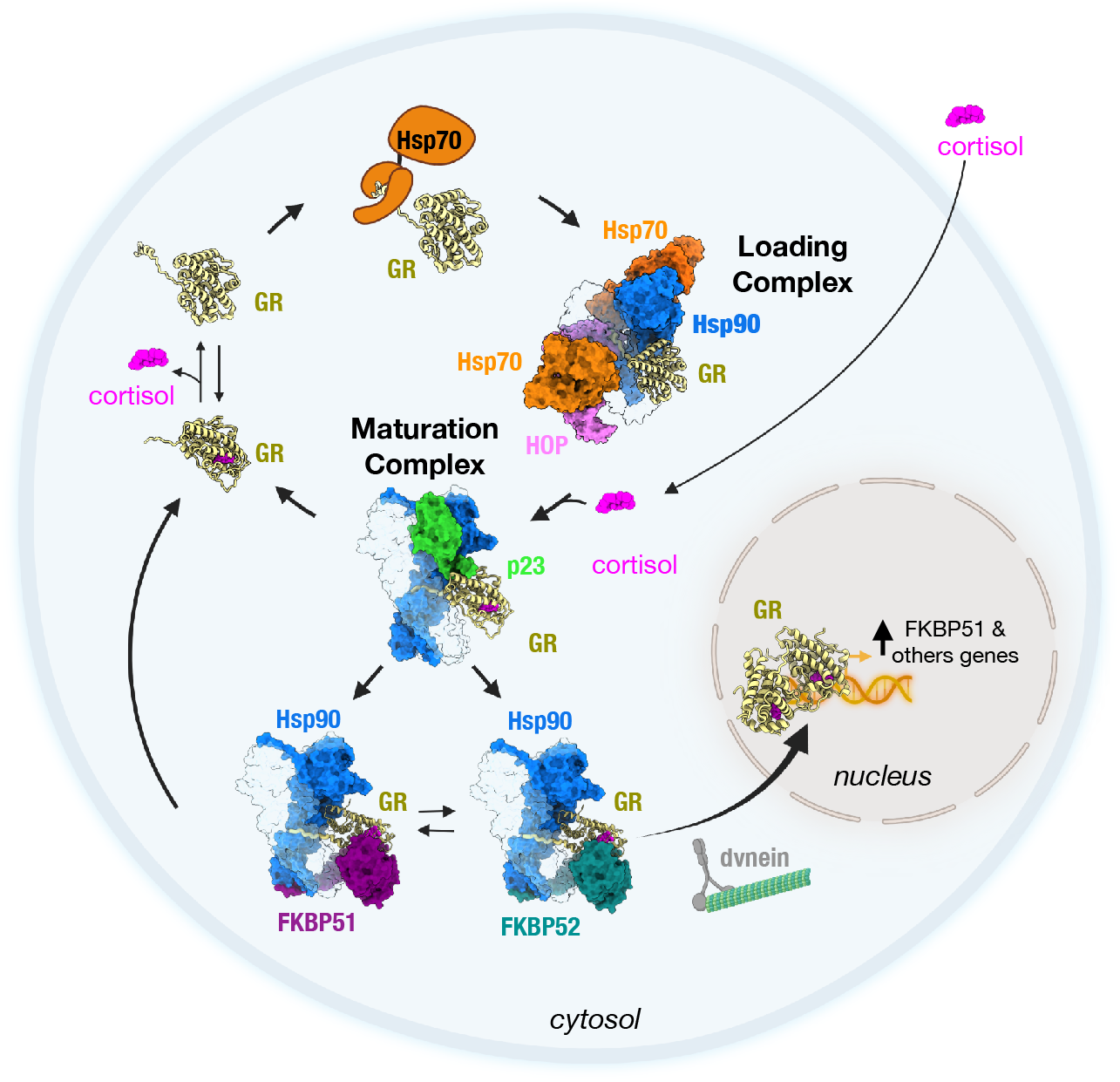
Mechanism of GR regulation by FKBP51 and FKBP52 in the GR chaperone cycle *in vivo*. Schematic of the GR chaperone cycle in the cell. Starting on the top left, GR (yellow, cartoon representation) is in dynamic equilibrium between cortisol-bound and unbound (apo) states. Hsp70 (orange) binds GR and locally unfolds GR to inhibit cortisol-binding, stabilizing GR in a partially unfolded, apo state. Hsp70 transfers the partially unfolded GR to Hsp90 (light and dark blue):Hop (pink) to form the GR-loading complex (Wang et al. 2022), in which GR is stabilized in a partially unfolded, apo state with the cortisol-binding pocket accessible. Cortisol (pink), which enters the cell through diffusion, binds to GR during the transition from the GR-loading complex to the GR-maturation complex when Hsp90 refolds the GR to a native conformation, sealing the cortisol-binding pocket through the refolding of the GR Helix 1 region (Noddings et al. 2022). In the GR-maturation complex, the cortisol-bound, folded GR is stabilized by Hsp90 and p23 (green), and is protected from Hsp70 re-binding. Depending on the relative concentrations of the FKBPs, either FKBP51 (purple) or FKBP52 (teal) can bind the GR:Hsp90:p23 complex, competing off p23, and stabilizing the rotated position of GR. FKBP51 sequesters GR:Hsp90 in the cytosol until ATP hydrolysis on Hsp90 allows release of GR back to the chaperone cycle. In contrast, FKBP52 promotes rapid nuclear translocation of GR:Hsp90 by acting as an adapter to the dynein/dynactin motor complex. Once in the nucleus, the cortisol-bound GR can dimerize, nucleate the assembly of transcriptional regulatory complexes, and activate the transcription of thousands of genes, including the gene for FKBP51 (*FKBP5*), leading to a negative feedback loop that regulates GR activity in the cell. The GR chaperone cycle also occurs in the absence of ligand and evidence supports preferential binding of FKBP51 over FKBP52 to apo GR:Hsp90 complexes, insuring the apo (inactivated) GR is not improperly translocated to the nucleus to regulate transcription.

Instead, we propose that ligand binds prior to the formation of either the GR-maturation complex or the GR:Hsp90:FKBP complexes, and that unbinding mostly occurs by recycling GR back to Hsp70, as previously described^4-6^.

After reaching the GR-maturation complex, the functional outcome for GR is dictated by FKBP51 and FKBP52, which compete to bind the GR:Hsp90 complex. FKBP52 stabilizes the ligand-bound GR, resulting in enhanced ligand affinity, and facilitates rapid GR nuclear translocation on dynein^22,24,25,62^, allowing GR to proceed with dimerization and activation of transcription in the nucleus. In contrast, FKBP51 binding keeps GR sequestered in the cytosol and recycles GR back to the chaperone cycle, inhibiting GR translocation and transcription activation. Interestingly, the expression of FKBP51, but not FKBP52, is upregulated by GR (as well as PR and AR), leading to a short negative feedback loop, which may help dampen chronic GR activation and signaling^27,63-67^. Thus, the relative concentrations of FKBP51 and FKBP52 in the cell dictate the level of GR activity *in vivo*^23,28,52^.

Beyond GR, FKBP51/52 are known to regulate the entire SHR class and given the sequence and structural conservation of the SHR LBDs at the FKBP contact sites, we propose FKBP51 and FKBP52 engage with the rest of the SHRs in a similar manner to GR (Extended Data Fig. 9a,b). Thus, FKBP51/52 can fine-tune the activity of these critical and clinically important signaling molecules and allow for crosstalk between the hormone signaling pathways. Altogether, we demonstrate how Hsp90 provides a platform for the FKBP co-chaperones to engage Hsp90 clients after Hsp90-dependent folding and promote the next step of client maturation, providing a critical layer of functional regulation.

## Acknowledgements

We thank members of the Agard Lab, past and present, including Ray Wang for the suggestion of this project and Elaine Kirschke for helpful discussions. We thank members of the José-Maria Carazo lab for continuous flexibility analysis on the cryo-EM datasets. We thank Jason Gestwicki for noting the potential importance of differential phosphorylation on the FKBPs. We thank David Bulkley, Glenn Gilbert, Zanlin Yu, and Eric Tse from the W.M. Keck Foundation Advanced Microscopy Laboratory at the University of California, San Francisco (UCSF) for EM facility maintenance and help with data collection. We also thank Matt Harrington and Joshua Baker-LePain for computational support with the UCSF Wynton cluster. C.M.N. is a National Cancer Institute Ruth L. Kirschstein Predoctoral Individual NRSA Fellow (F31CA265084-02). The work was supported by NIH grants R35GM118099 (D.A.A.), S10OD020054 (D.A.A.), S10OD021741 (D.A.A.), P20GM104420 (J.L.J.), and R01GM127675 (J.L.J.).

## Author Contributions

C.M.N. designed and executed biochemical experiments, cryo-EM sample preparation, data collection, data processing, and model building. J.L.J. executed yeast *in vivo* assays and interpreted the results. C.M.N. and D.A.A. conceived the project, interpreted the results, and wrote the manuscript.

## Competing Interests

The authors declare no competing interests.

## Notes

### Competing Interest Statement

The authors have declared no competing interest.

### Summary of Updates

Figure 4a,b was mislabeled as "FKBP52", which should be "FKBP51"

