## Supplementary Figures and Methods for "Cryo-EM reveals how Hsp90 and FKBP immunophilins co-regulate the Glucocorticoid Receptor"

This PDF file includes:

Extended Data Figures 1 to 9; Extended Data Table 1

Guide to Supplementary Movies 1 to 6

Materials and Methods

References for Materials and Methods

#### Extended Data Figure 1

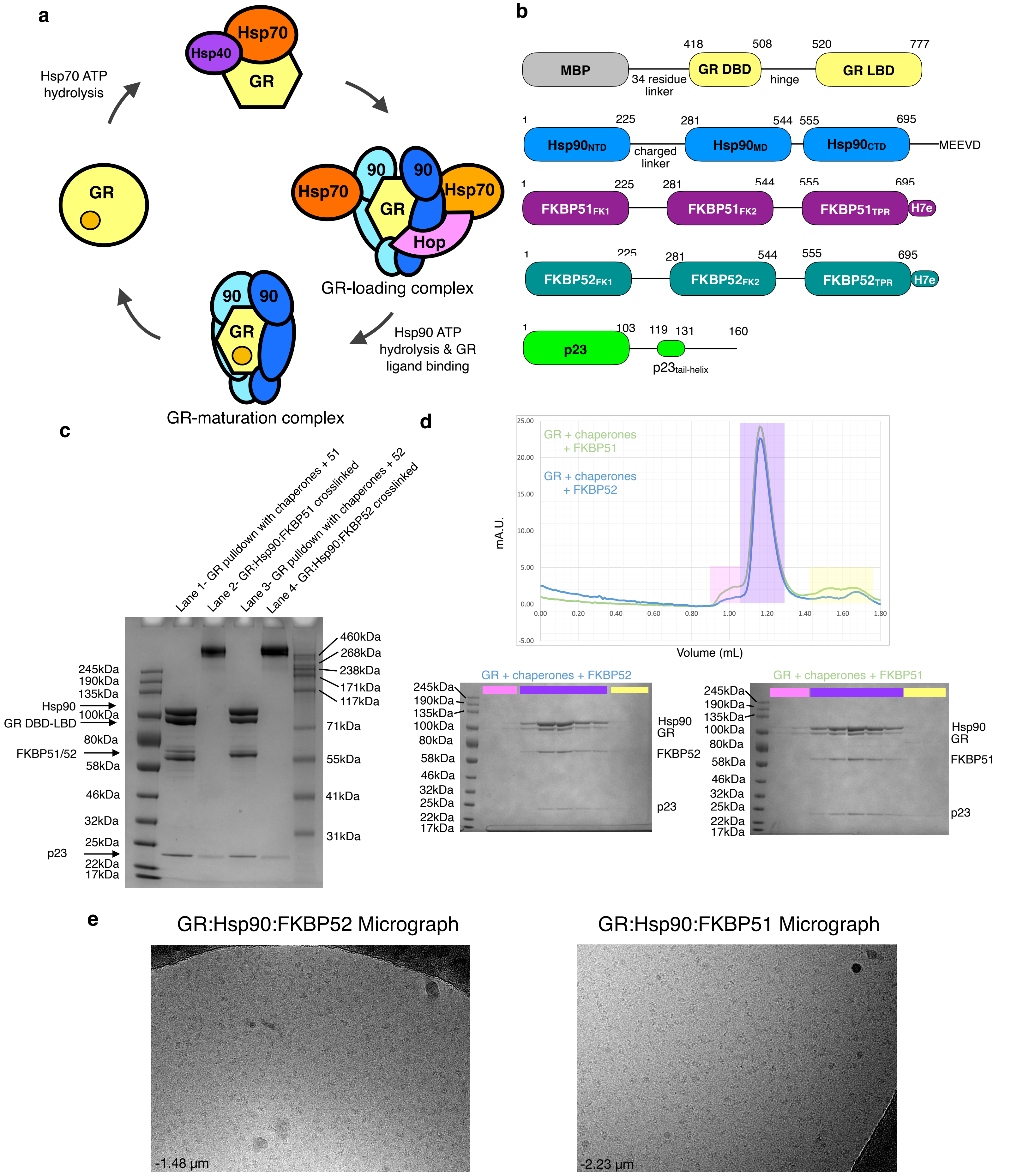

**Extended Data Fig. 1 | Sample Preparation**
**a,** The *in vitro* reconstituted GR-chaperone cycle. On the left, GR alone is active and able to bind ligand. Hsp70, aided by the co-chaperone Hsp40, engages GR and inhibits ligand binding. Hsp70 loads GR onto Hsp90 and Hop, which forms the "GR-loading complex". GR is stabilized in an inhibited, partially unfolded conformation by semi-closed Hsp90, Hsp70, and Hop (PDB ID 7KW7). Hsp70 and Hop are released, Hsp90 hydrolyzes ATP to fully close, and the co-chaperone p23 binds, forming the "GR-maturation complex" (PDB ID 7KRJ). GR binds ligand in the transition from the GR-loading complex to the GR-maturation complex. In the maturation complex, GR is in a fully folded, native conformation and bound to ligand. Upon Hsp90 re-opening, GR is released from the complex to return to the cycle. **b,** Domain organization of the proteins in the GR:Hsp90:FKBP complexes as well as p23. **c,** Coomassie-stained SDS-PAGE (4-12% acrylamide gel) with elution from the MBP-GR pulldown from the *in vitro* reconstituted GR chaperone cycle. Lanes 1 and 2 show the elution from the FKBP51-containing sample, while Lanes 3 and 4 show the elution from the FKBP52-containing sample. Assay conditions are as follows- Lane 1: 5 mM MBP-GR, 2 mM Hsp40, 5 mM Hsp70, 5 mM Hop, 15 mM Hsp90, 15 mM Bag-1, 30 mM FKBP51, 5 mM ATP, 20mM Sodium Molybdate; Lane 2: sample from Lane 1 after size exclusion chromatography (**d**) and chemical crosslinking with 0.02% glutaraldehyde; Lane 3: 5 mM MBP-GR, 2 mM Hsp40, 5 mM Hsp70, 5 mM Hop, 15 mM Hsp90, 15 mM Bag-1, 30 mM FKBP52, 5 mM ATP, 20mM Sodium Molybdate; Lane 4: sample from Lane 3 after size exclusion chromatography (**d**) and chemical crosslinking with 0.02% glutaraldehyde. **d,** S200 3.2/300 size exclusion chromatography (SEC) profile of the elution from the MBP-GR pulldown. The green trace represents the SEC profile of the from the reconstituted GR chaperone cycle with FKBP51, while the blue trace represents the SEC profile from the reconstituted GR chaperone cycle with FKBP52. mAU=milli-absorbance units. Coomassie-stained SDS-PAGE (4-12% acrylamide gel) of the fractions from size exclusion chromatography corresponding to the GR:Hsp90:FKBP52 sample (left) or the GR:Hsp90:FKBP51 sample (right). Colors indicate which gel lanes correspond to specific regions of the size exclusion chromatography profile. Sample fractions from the region highlighted in purple were collected and used for cryo-EM data collection. This experiment was repeated 11 independent times with similar results. **e,** Representative electron micrograph for the cryo-EM dataset of the GR:Hsp90:FBKP52 complex (left) (-1.48 mm defocus) and GR:Hsp90:FKBP51 complex (right) (-2.23 mm defocus). A total of 11,162 and 26,413 micrographs were obtained, respectively.

### Extended Data Figure 2

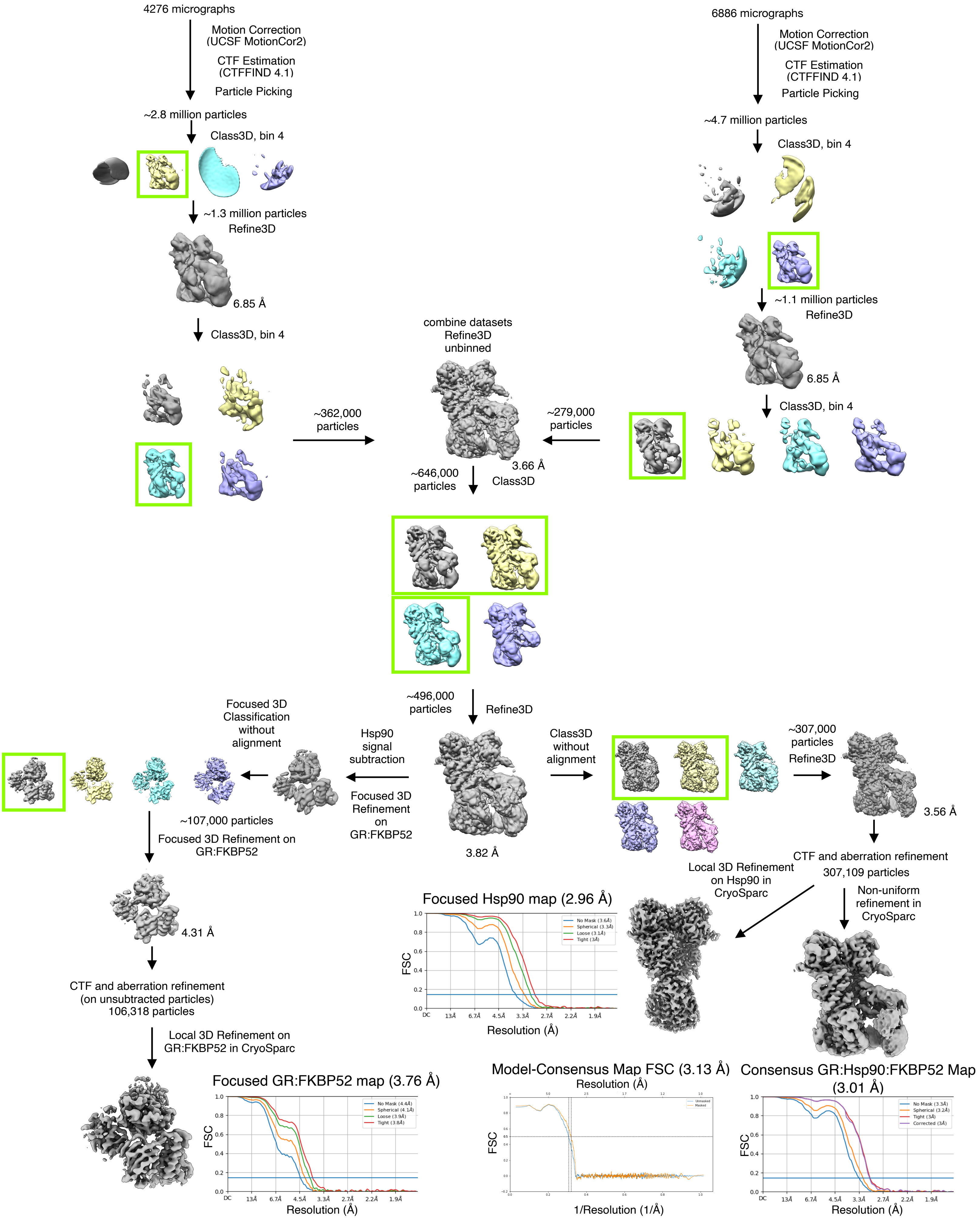

**Extended Data Fig. 2 | Cryo-EM Data Analysis for the GR:Hsp90:FKBP52 Complex**
Cryo-EM data processing procedure for the GR:Hsp90:FKBP52 complex performed in RELION and CryoSparc. Gold-standard Fourier shell correlation (GSFSC) curves of the final 3D reconstructions, including the focused maps and the consensus map, are shown (bottom). The blue lines intercept the y-axis at an FSC value of 0.143. Map-to-model FSC curves between the GR:Hsp90:FKBP52 atomic model and the consensus GR:Hsp90:FKBP52 map. The black dotted line intercepts the y-axis at an FSC value of 0.5.

**Extended Data Figure 3**

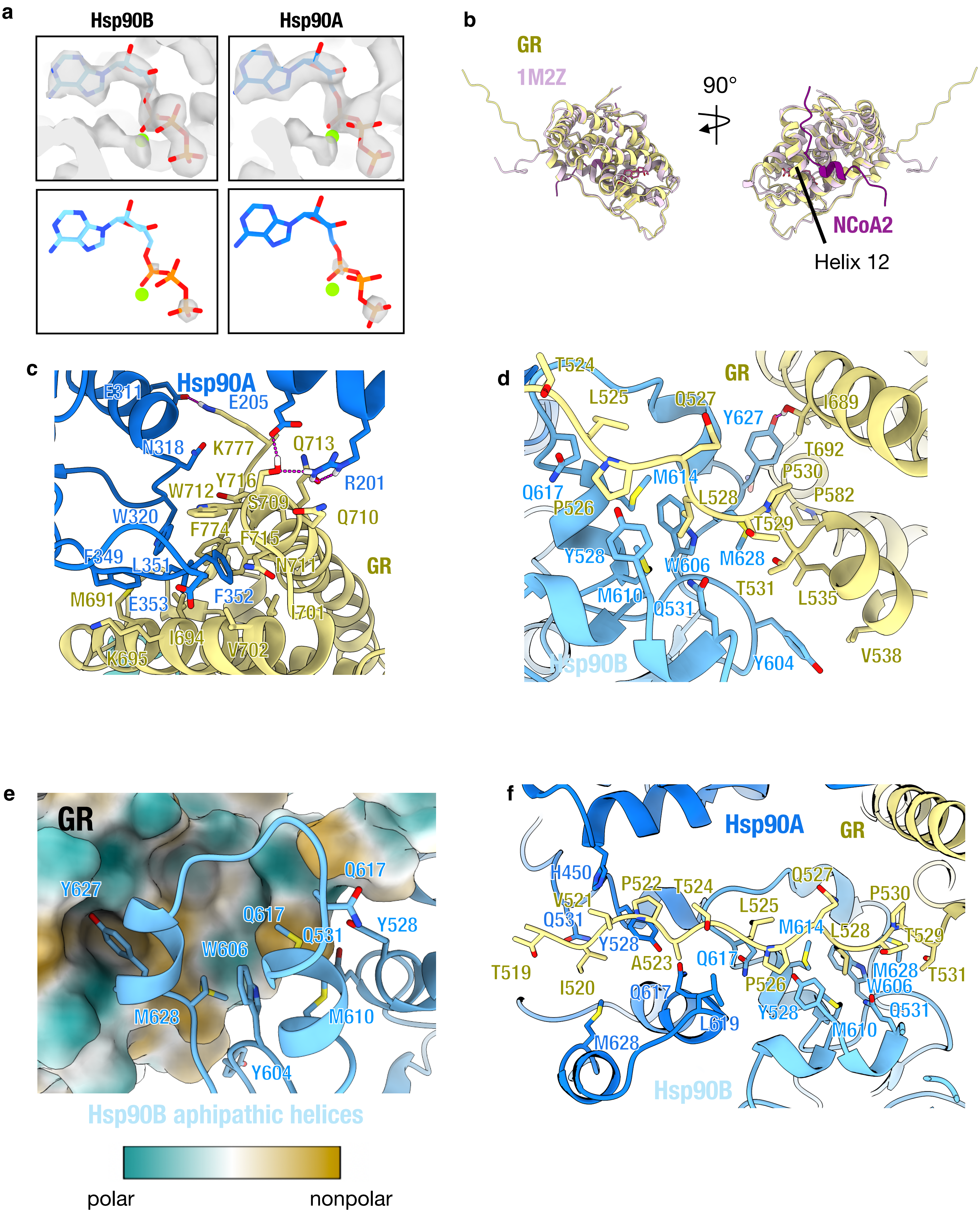

**Extended Data Fig. 3 | Hsp90:GR Interfaces in the GR:Hsp90:FKBP52 Complex**
**a,** Hsp90:GR:FKBP52 complex map density with atomic model showing ATP-magnesium density in both Hsp90 protomers (Hsp90A/B). Bottom images show increased contour level on the map density to indicate that the ATP g-phosphate position has much stronger density relative to the a and b-phosphates, likely corresponding to molybdate, which may act as a g-phosphate analog (see **Methods**). **b,** Atomic model of GR from the GR:Hsp90:FKBP52 complex (yellow) compared with GR from the crystal structure (PDB ID 1M2Z) (light pink) with co-activator peptide NCoA2 (purple) and ligand (dark pink). $GR_{Helix\ 12}$ is indicated. **c-f,** Atomic model of GR:Hsp90:FKBP52 complex with Hsp90A (dark blue), Hsp90B (light blue), GR (yellow). Side chains in contact between GR and Hsp90 are shown, along with hydrogen bonds (dashed pink lines).**c,** Interface 1 of the GR:Hsp90 interaction depicting the GR hydrophobic patch (GR Helices 9 and 10) interacting with the Hsp90A Src loop ($Hsp90^{345-360}$), $Hsp90A^{W320}$, and Hsp90A NTD/MD helices. **d,** Interface 2 of the GR:Hsp90 interaction depicting GR pre-Helix 1 strand and Helix 1 packing up against the Hsp90B amphipathic a-helices. **e,** Interface 2 of the GR:Hsp90 interaction depicting GR in surface representation colored by hydrophobicity (green = polar, brown = nonpolar) with $Hsp90B^{Y627}$ sticking into a the BF3 druggable hydrophobic pocket. **f,** Interface 3 of the GR:Hsp90 interaction depicting the GR pre-Helix 1 strand threading through the Hsp90 lumen between Hsp90A and Hsp90B.

**Extended Data Figure 4**

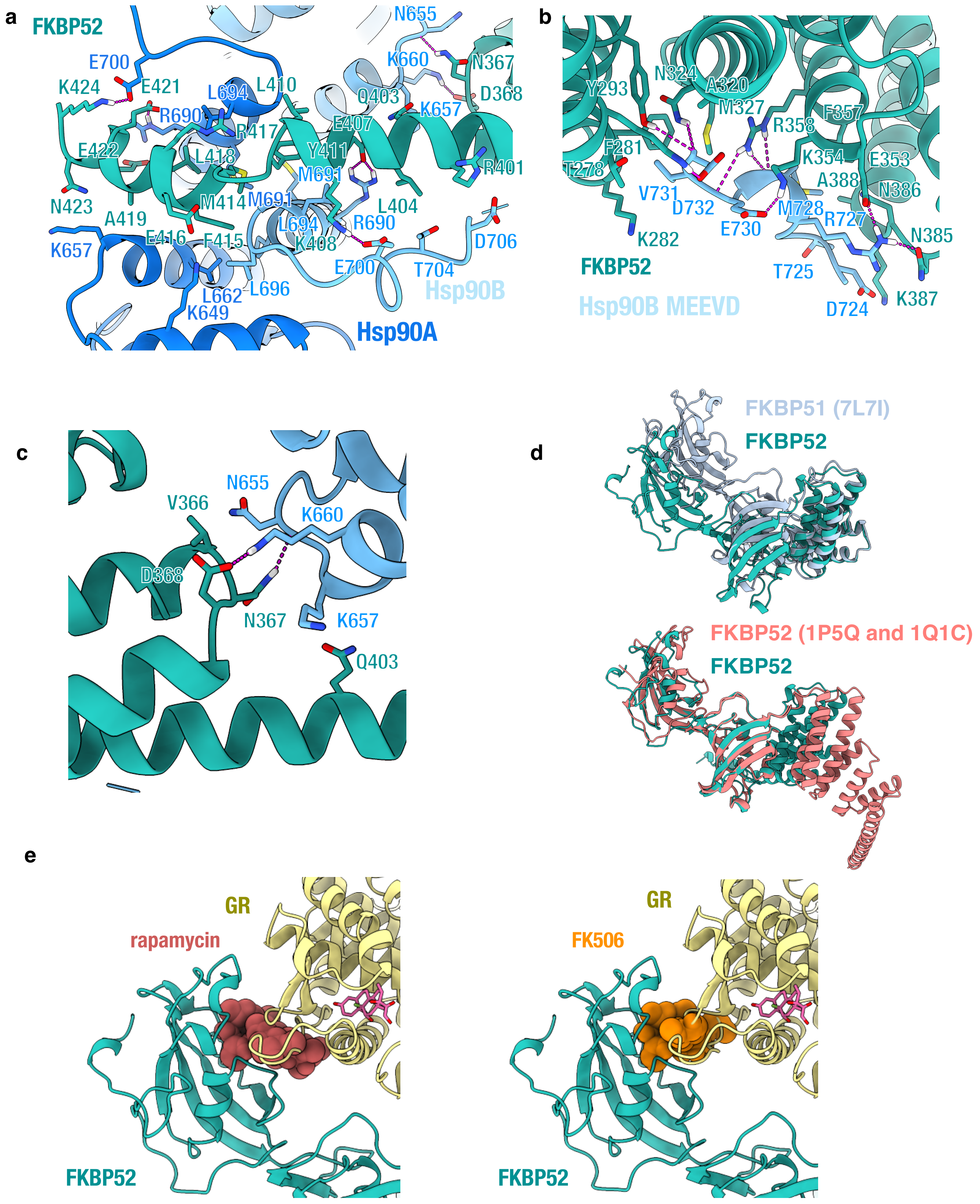

**Extended Data Fig. 4 | Hsp90:FKBP52 Interfaces in the GR:Hsp90:FKBP52 Complex**
Atomic model of the GR:Hsp90:FKBP52 complex with Hsp90A (dark blue), Hsp90B (light blue), GR (yellow), and FKBP52 (teal). Side chains in contact between Hsp90 and FKBP52 are shown, along with hydrogen bonds (dashed pink lines). **a,** Interface 1 of the Hsp90:FKBP52 interaction depicting the FKBP52 TPR H7e binding to the Hsp90A/B CTD dimer interface. The helix of FKBP52 H7e breaks to fit into the cleft formed by the Hsp90 CTDs. **b,** Interface 2 of the Hsp90:FKBP52 interaction depicting the Hsp90B MEEVD motif binding the FKBP52 TPR helical bundle. **c,** Interface 3 of the Hsp90:FKBP52 interaction depicting the FKBP52 TPR helices 5 and 6 binding to the Hsp90B CTD. **d,** FKBP52 (teal) from the GR:Hsp90:FKBP52 atomic model aligned with the cryo-EM structure of FKBP51 (light blue) (PDB ID 7L7I) (top) and crystal structures of FKBP52 (PDB ID 1P5Q, 1Q1C) (bottom) showing the difference in interdomain angles. 1P5Q contains the FKBP52 FK1 and FK2 domain, while 1Q1C contains the FKBP52 FK2 and TPR domains. **e,** The GR:Hsp90:FKBP52 atomic model with FKBP52 (teal), GR (yellow), and dexamethasone (pink) with proline-isomerase inhibitors, rapamycin (brown) or FK506 (orange), docked into the atomic model to indicate the steric clash with GR. Rapamycin was docked in based on the FKBP52:rapamycin crystal structure (PDB ID 4DRJ) and FK506 was docked in based on the FKBP52:FK506:FRB crystal structure (PDB ID 4LAX).

**Extended Data Figure 5**

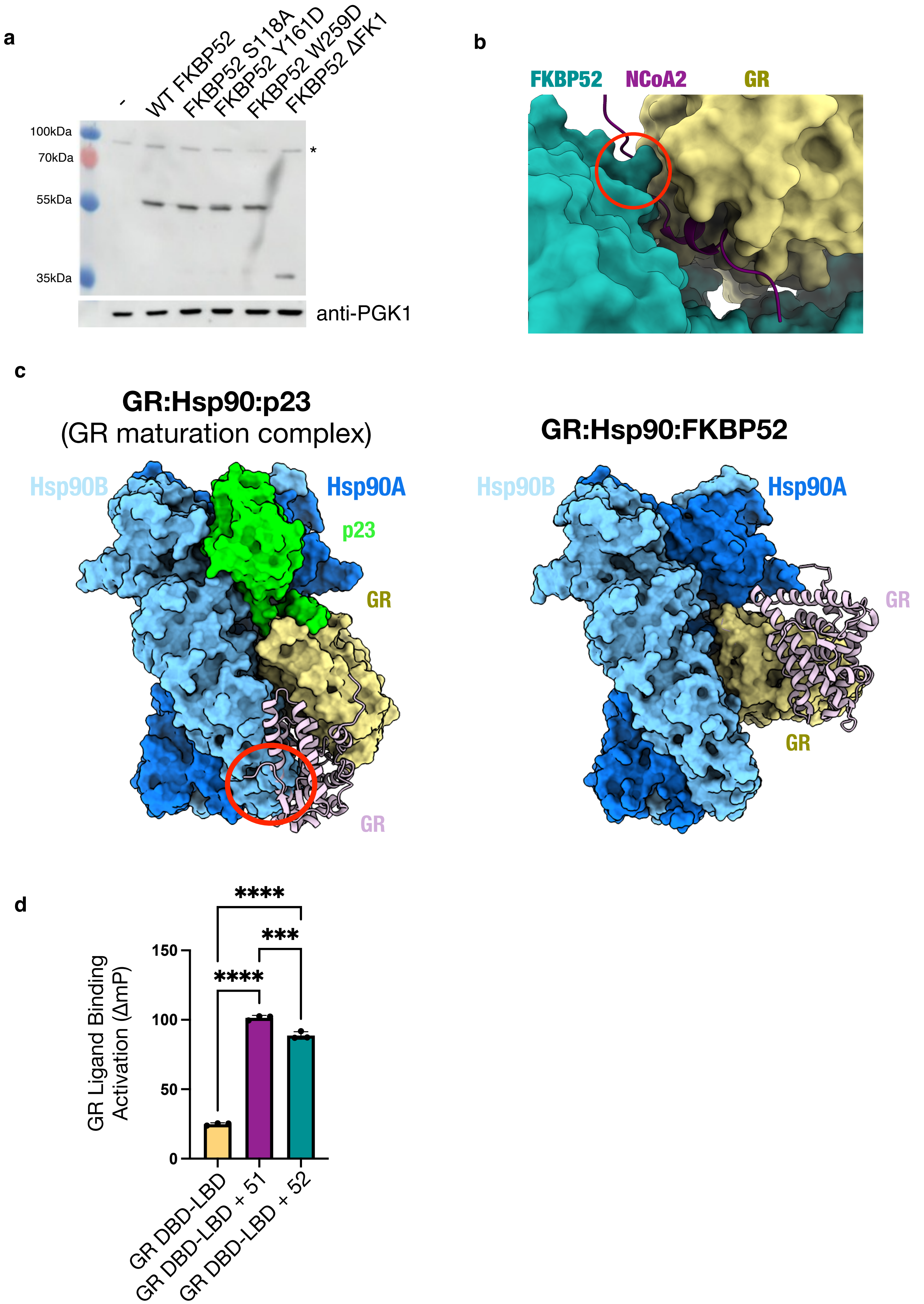

**Extended Data Fig. 5 | Analysis of the GR:Hsp90:FKBP52 Structure**
**a,** Expression of human FKBP52 or FKBP52 mutants in wild-type yeast strain JJ762 assayed by immunoblot with a monoclonal antibody specific for FKBP52. An antibody against PGK1 was used as a loading control. The asterisk marks an unknown protein that cross reacts with the anti-FKBP52 antibody. **b,** Atomic model of the GR:Hsp90:FKBP52 complex shown in surface representation. FKBP52 (teal), GR (yellow). The NCoA2 (nuclear coactivator 2) co-activator peptide is docked in based on the GR:NCoA2 crystal structure (PDB ID 1M2Z). While most of the coactivator peptide binding is sterically permitted, the N-terminus of NCoA2 clashes with the FKBP52 TPR domain (red circle). **c,** Atomic models of the GR-maturation complex (GR:Hsp90:p23) (left) and the GR:Hsp90:FKBP52 complex without FKBP52 (right) depicting that GR LBD dimerization is permitted once FKBP52 is released. Hsp90A (dark blue, surface representation), Hsp90B (light blue, surface representation), GR (yellow, surface representation), p23 (green, surface representation). In both complexes, the GR LBD dimerization site is accessible, however; binding of the second GR LBD (light pink) to the GR-maturation complex clashes with the Hsp90B CTD, shown with a red circle (left). Binding of the second GR LBD (light pink) to the GR:Hsp90:FKBP52 complex (right). Docking of the dimerized GR LBD is based on the GR LBD dimer crystal structure (PDB ID 1M2Z). **d,** Equilibrium binding of 10 nM fluorescent dexamethasone to 100 nM GR DBD-LBD with addition of 15 mM FKBP51 (“51”) or FKBP52 (“52”) measured by fluorescence polarization (mean±SD). n=3 biologically independent samples per condition. Fluorescence polarization values are baseline subtracted in accordance with the measured fluorescent dexamethasone baseline polarization value. Statistical significance was evaluated by an ordinary one-way ANOVA ($F_{(2,6)} = 1414$, $p < 0.0001$) with *post-hoc* Tukey’s multiple comparisons test. P-values: p(GR vs. GR + 51) < 0.0001, p(GR vs. GR + 52) < 0.0001, p(GR + 51 vs. GR + 52) = 0.004. (n.s. P ≥ 0.05; * P ≤ 0.05; ** P ≤ 0.01; *** P ≤ 0.001; **** P ≤ 0.0001).

### Extended Data Figure 6

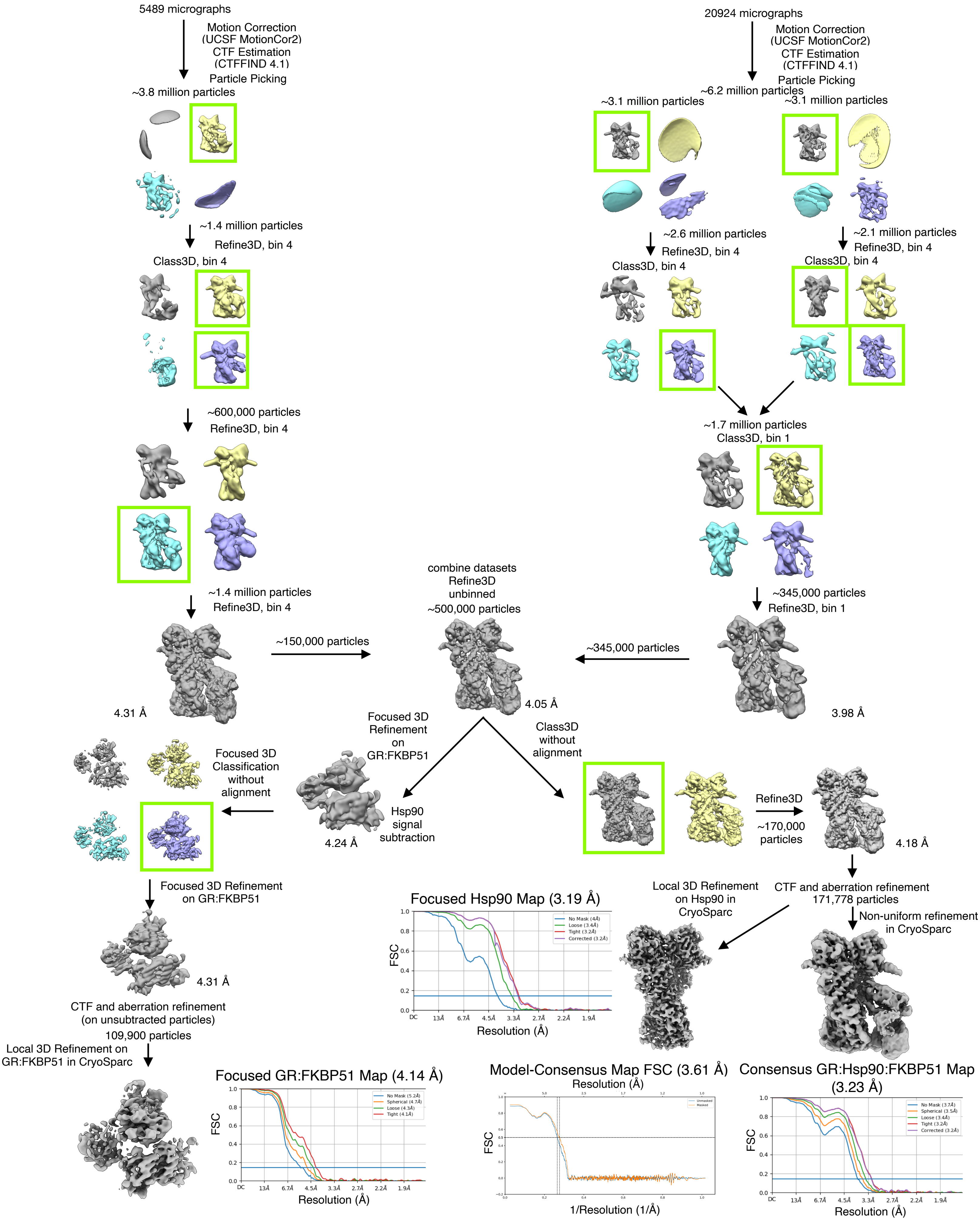

**Extended Data Fig. 6 | Cryo-EM Data Analysis for the GR:Hsp90:FKBP51 Complex**
Cryo-EM data processing procedure for the GR:Hsp90:FKBP51 complex performed in RELION and CryoSparc. Gold-standard Fourier shell correlation (GSFSC) curves of the final 3D reconstructions, including the focused maps and the consensus map, are shown (bottom). The blue lines intercept the y-axis at an FSC value of 0.143. Map-to-model FSC curves between the GR:Hsp90:FKBP51 atomic model and the consensus GR:Hsp90:FKBP51 map. The black dotted line intercepts the y-axis at an FSC value of 0.5.

Extended Data Figure 7

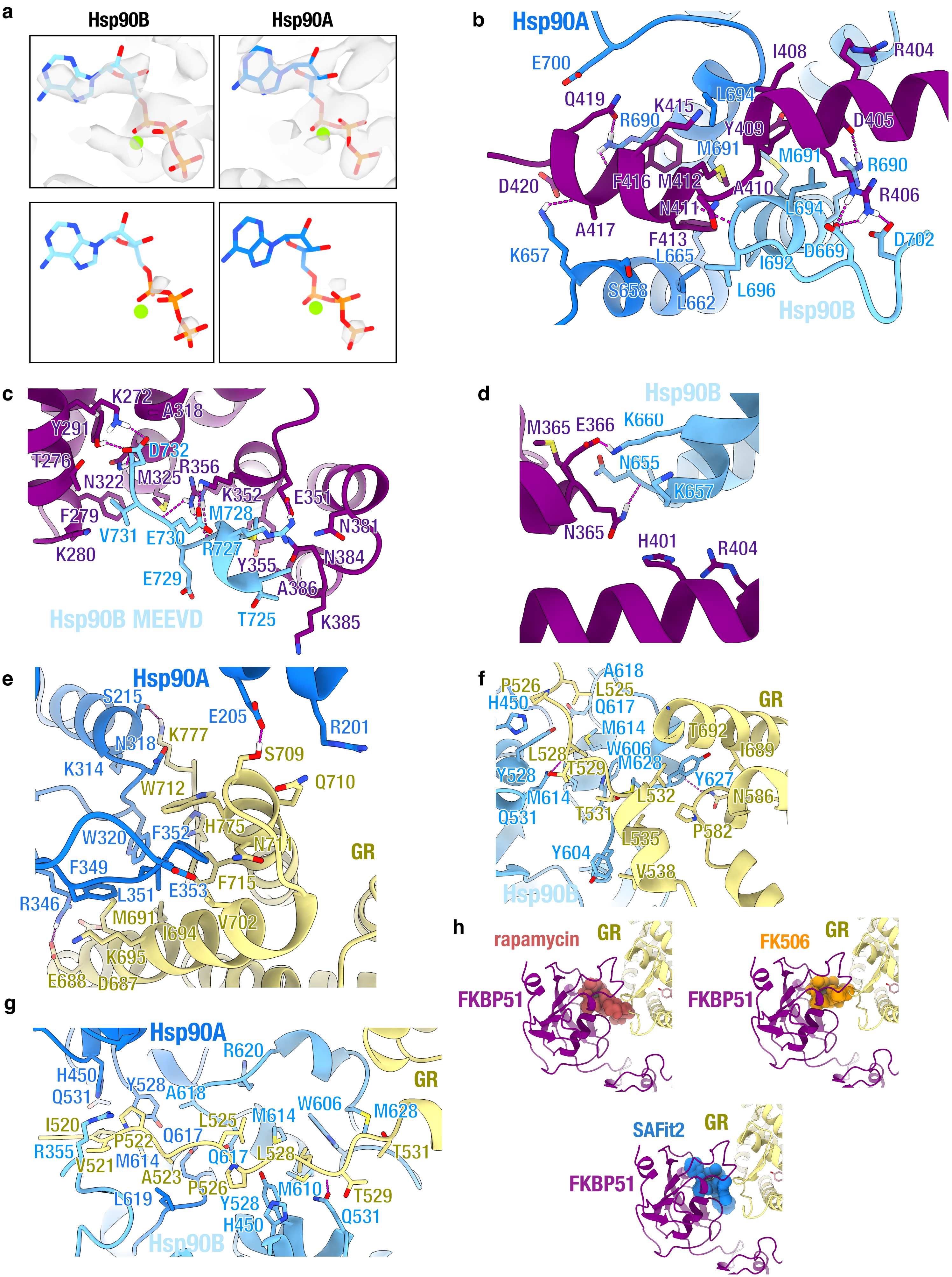

**Extended Data Fig. 7 | Hsp90:FKBP51 Interfaces in the GR:Hsp90:FKBP51 Complex**
Atomic model of the GR:Hsp90:FKBP51 complex with Hsp90A (dark blue), Hsp90B (light blue), GR (yellow), and FKBP51 (purple). Side chains in contact between Hsp90 and FKBP51 or Hsp90 and GR are shown, along with hydrogen bonds (dashed pink lines). **a,** Hsp90:GR:FKBP51 complex map density with atomic model showing ATP-magnesium density in both Hsp90 protomers (Hsp90A/B). Bottom images show increased contour level on the map density to indicate that the ATP g-phosphate position has much stronger density relative to the a and b-phosphates, likely corresponding to molybdate, which may act as a g-phosphate analog (see **Methods**). **b,** Interface 1 of the Hsp90:FKBP51 interaction depicting the FKBP51 TPR H7e binding to the Hsp90A/B CTD dimer interface. The helix of FKBP51 H7e breaks to fit into the cleft formed by the Hsp90 CTDs. **c,** Interface 2 of the Hsp90:FKBP51 interaction depicting the Hsp90B MEEVD motif binding the FKBP51 TPR helical bundle **d,** Interface 3 of the Hsp90:FKBP51 interaction depicting the FKBP51 TPR helices 5 and 6 binding to the Hsp90B CTD. **e,** Interface 1 of the GR:Hsp90 interaction depicting the GR hydrophobic patch (GR Helices 9 and 10) interacting with the Hsp90A Src loop ($Hsp90^{345\text{-}360}$), $Hsp90A^{W320}$, and Hsp90A NTD/MD helices. **f,** Interface 2 of the GR:Hsp90 interaction depicting GR pre-Helix 1 strand and Helix 1 packing up against the Hsp90B amphipathic a-helices. **g,** Interface 3 of the GR:Hsp90 interaction depicting the GR pre-Helix 1 strand threading through the Hsp90 lumen between Hsp90A and Hsp90B. **h,** The GR:Hsp90:FKBP51 atomic model with FKBP51 (purple), GR (yellow), and dexamethasone (pink) with proline-isomerase inhibitors, rapamycin (brown) or FK506 (orange) docked into the atomic model to indicate the steric clash with GR. Rapamycin was docked in based on the FKBP52:rapamycin crystal structure (PDB ID 4DRI) and FK506 was docked in based on the FKBP52:FK506:FRB crystal structure (PDB ID 3O5R). The FKBP52-specific inhibitor SAFit2 was docked into the atomic model to indicate there is no steric clash with GR at the backbone level (although some side chains clash). SAFit2 was docked in based on the FKBP51:SAFit2 crystal structure (PDB ID 6TXX).

**Extended Data Figure 8**

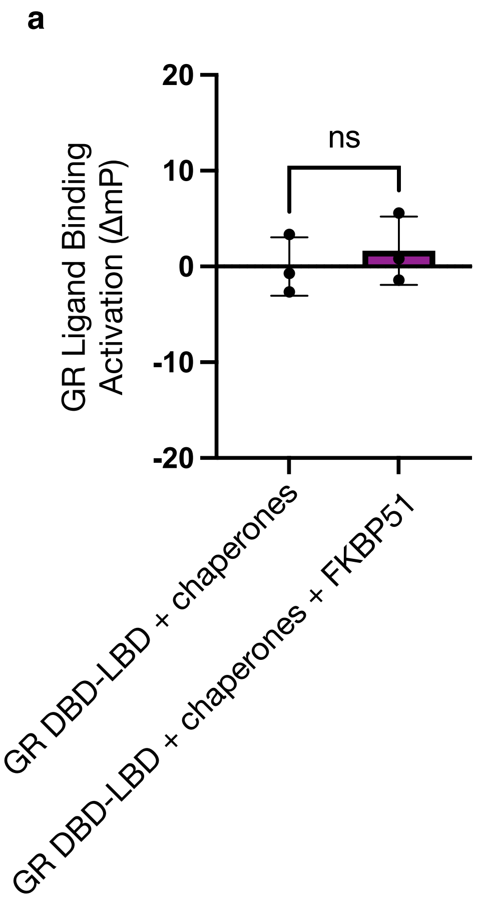


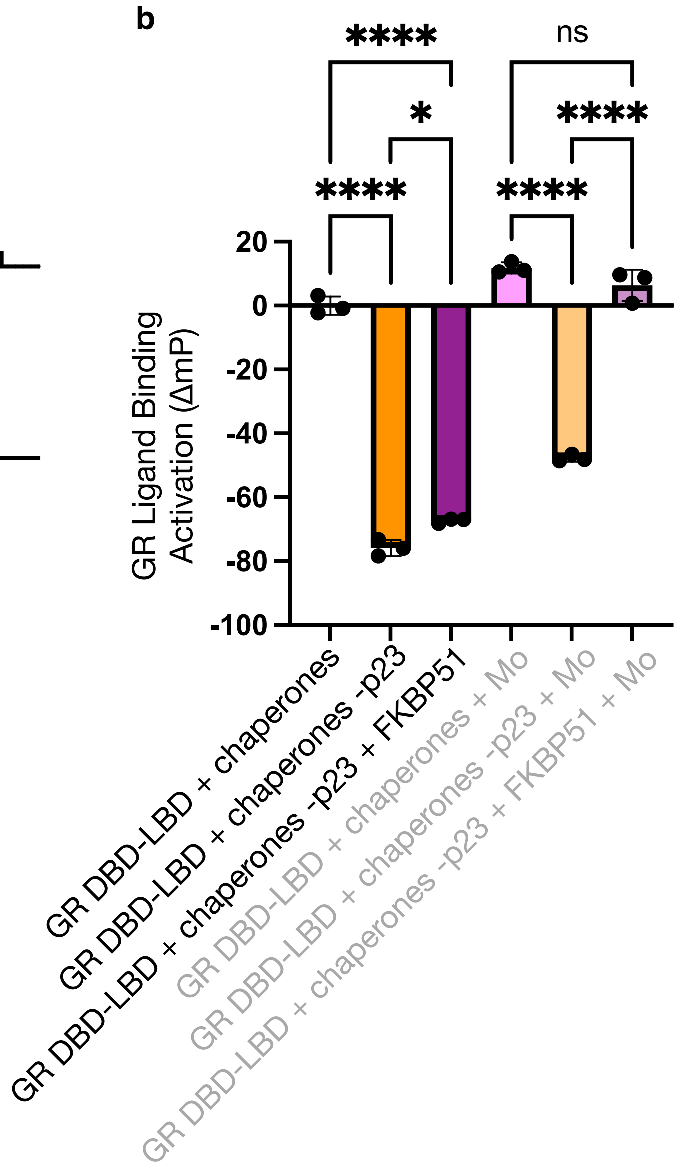

**Extended Data Fig. 8 | Effect of FKBP51 on GR Ligand Binding *in vitro***
**a,** Equilibrium binding of 10nM fluorescent dexamethasone to 100nM GR DBD-LBD with chaperones and FKBP51. “Chaperones”= 15uM Hsp70, Hsp90, Hop, p23, and FKBP51; 2uM Ydj1 and Bag-1. Statistical significance was evaluated by an unpaired two-tailed t test, p-value =0.5737. (n.s. P ≥ 0.05; * P ≤ 0.05; ** P ≤ 0.01; *** P ≤ 0.001; **** P ≤ 0.0001.**b,** Equilibrium binding of 10nM fluorescent dexamethasone to 100nM GR DBD-LBD with chaperones, FKBP51, and sodium molybdate (“Mo”). “Chaperones”= 15uM Hsp70, Hsp90, Hop, p23, and FKBP51; 2uM Ydj1 and Bag-1. Statistical significance was evaluated by an ordinary one-way ANOVA ($F_{(6,12)}$ = 647.1, p < 0.0001) with *post-hoc* Šídák’s multiple comparisons test. P-values: p(Chaperones vs. Chaperones -p23) < 0.0001, p(Chaperones vs. Chaperones -p23 + 51) < 0.0001= 0.0287, p(Chaperones -p23 vs. Chaperones -p23 + 51) = 0.0123, p(Chaperones + Mo. Vs. Chaperones -p23 + Mo. ) < 0.0001, p(Chaperones + Mo. Vs. Chaperones -p23 + 51 + Mo. ) = 0.1640, p(Chaperones -p23 + Mo. Vs. Chaperones -p23 + 51 + Mo. ) < 0.0001. (n.s. P ≥ 0.05; * P ≤ 0.05; ** P ≤ 0.01; *** P ≤ 0.001; **** P ≤ 0.0001).

**Extended Data Figure 9**

**a**

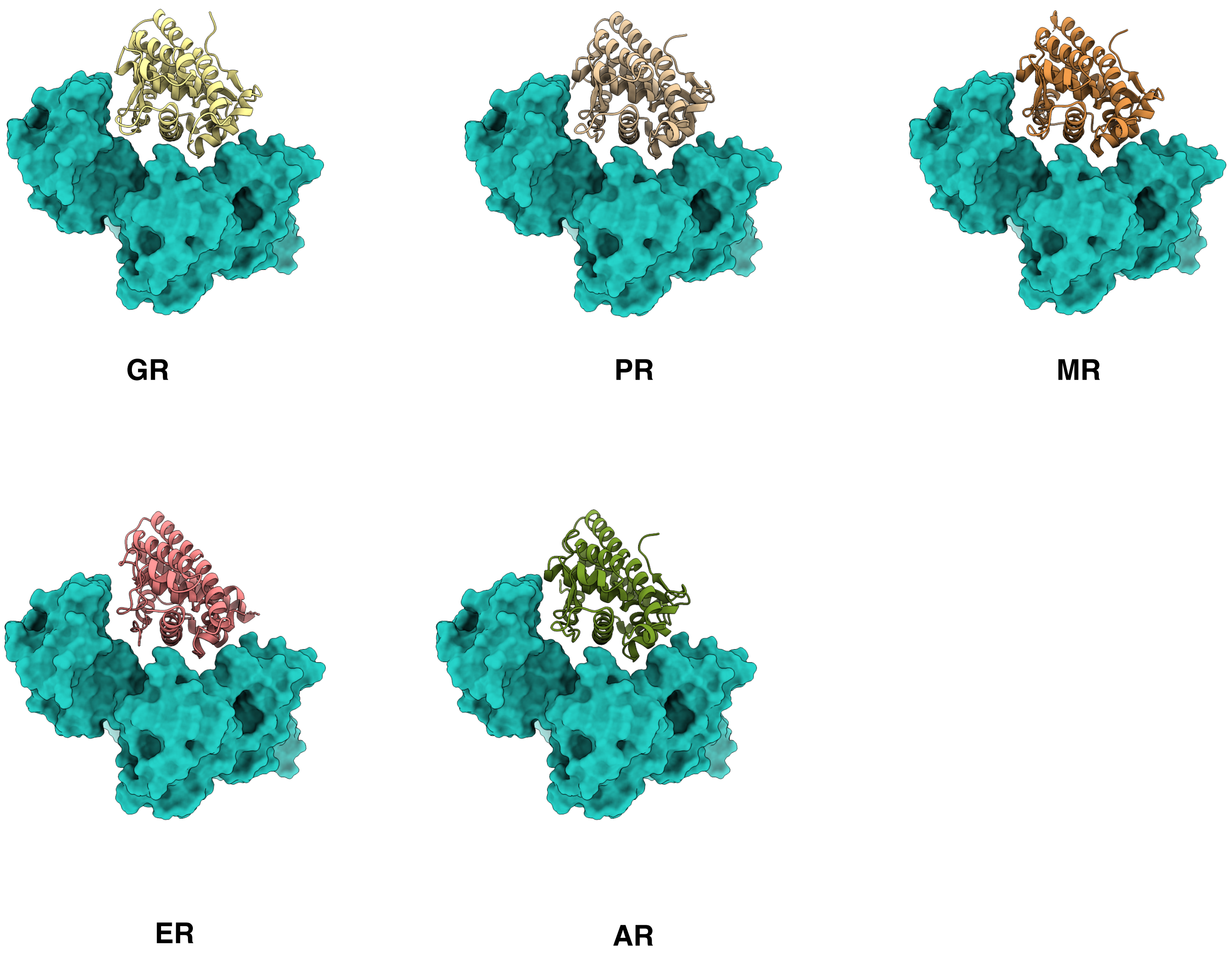


**b**

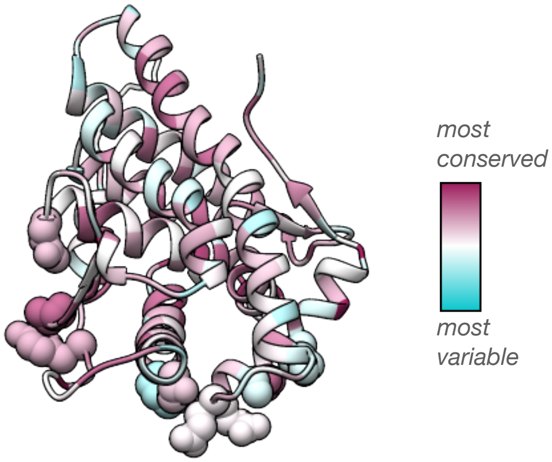

**Extended Data Fig. 9 | Modeling FKBP Binding to All Five Steroid Hormone Receptors (SHRs)**
**a,** FKBP52 bound to the glucocorticoid receptor (GR, yellow), progesterone receptor (PR, tan), mineralocorticoid receptor (MR, orange), estrogen receptor (ERa, pink), or androgen receptor (AR, green) based on the structure of the GR:Hsp90:FKBP52 complex. Due to the structural conservation of the LBDs across the five SHRs, all SHRs fit well with FKBP52 from the GR:Hsp90:FKBP52 atomic model, with no backbone clashes between the SHRs and FKBP52. The PDB IDs used to dock in the SHRs are as follows: PR (1A28), MR (2AA7), ERa (1ERE), AR (1T7R). **b,** Sequence conservation across human steroid hormone receptors (GR, MR, ERa, ERb, AR) plotted onto the GR structure from the GR:Hsp90:FKBP52 atomic model. Residues are colored from most variable (cyan) to most conserved (maroon).

**Extended Data Table 1 | Cryo-EM data collection, refinement and validation statistics**

| | GR:Hsp90:FKBP52 (EMDB-29068) (PDB 8FFV) | | GR:Hsp90:FKBP51 (EMDB-29069) (PDB 8FFW) | |
|---|---|---|---|---|
| **Data collection and processing** | Dataset I | Dataset II | Dataset III | Dataset IV |
| Magnification | 105,000 | 105,000 | 105,000 | 105,000 |
| Voltage (kV) | 300 | 300 | 300 | 300 |
| Electron exposure (e–/Å$^2$) | 69 | 69 | 69 | 45.8 |
| Defocus range (μm) | 0.8-2.0 | 0.8-2.0 | 0.8-2.0 | 0.8-2.0 |
| Pixel size (Å) | 0.835 | 0.835 | 0.835 | 0.835 |
| Symmetry imposed | C1 | C1 | C1 | C1 |
| Initial particle images (no.) | 4,736387 | 2,783,284 | 3,788,005 | 6,268,573 |
| Consensus Map resolution (Å) | 3.01 | | 3.23 | |
| FSC threshold | 0.143 | | 0.143 | |
| Final particle images (no.) | 307,109 | | 171,778 | |
| Hsp90 Focused Map resolution (Å) | 2.96 | | 3.19 | |
| FSC threshold | 0.143 | | 0.143 | |
| Final particle images (no.) | 307,109 | | 171,778 | |
| GR:FKBP Focused map resolution (Å) | 3.76 | | 4.14 | |
| FSC threshold | 0.143 | | 0.143 | |
| Final particle images (no.) | 106,318 | | 109,900 | |
| **Refinement** | | | | |
| Initial model used (PDB code) | 7KRJ, AF-Q02790, 5NJX | | 7KRJ, 7L7I, 5NJX | |
| Model resolution (Å) | 3.13 | | 3.61 | |
| FSC threshold | 0.5 | | 0.5 | |
| Map sharpening *B* factor (Å$^2$) | -30 | | -30 | |
| Model composition | | | | |
| Non-hydrogen atoms | 15829 | | 15756 | |
| Protein residue atoms | 15737 | | 15664 | |
| Ligand atoms | 92 | | 92 | |
| Mean *B* factors [min-max] (Å$^2$) | | | | |
| Protein | 209.15 [0.03-600.0] | | 244.41 [60.07-600.0] | |
| Ligand | 140.07 [15.18-409.30] | | 222.63 [75.96-528.72] | |
| R.m.s. deviations | | | | |
| Bond lengths (Å) | 0.023 | | 0.024 | |
| Bond angles (°) | 1.837 | | 1.936 | |
| Validation | | | | |
| MolProbity score | 0.81 | | 0.84 | |
| Clashscore | 1.07 | | 1.24 | |
| Poor rotamers (%) | 0.00 | | 0.00 | |
| Ramachandran plot | | | | |
| Favored (%) | 98.70 | | 98.55 | |
| Allowed (%) | 1.19 | | 1.40 | |
| Disallowed (%) | 0.10 | | 0.05 | |

#### Guide to Supplementary Movies

**Supplementary Movie 1**

CryoSparc 3D variability analysis component 0 (side view) showing the GR:Hsp90:FKBP52 focused map density and atomic model. The FKBP52 (teal) FK1 domain displays continuous motion and is dynamically associated with GR (yellow) relative to the other FKBP52 domains (FK2, TPR).

**Supplementary Movie 2**

CryoSparc 3D variability analysis component 2 showing the GR:Hsp90:FKBP52 focused map density and atomic model. The FKBP52 (teal) FK1 domain displays continuous motion and is dynamically associated with GR (yellow) relative to the other FKBP52 domains (FK2, TPR).

**Supplementary Movie 3**

CryoSparc 3D variability analysis component 0 (top view) showing the GR:Hsp90:FKBP52 focused map density and atomic model. The FKBP52 (teal) proline-rich loop on the FK1 domain displays continuous motion and is dynamically associated with GR (yellow).

**Supplementary Movie 4**

CryoSparc 3D variability analysis component 2 (top view) showing the GR:Hsp90:FKBP52 focused map density and atomic model. The FKBP52 (teal) proline-rich loop on the FK1 domain displays continuous motion and is dynamically associated with GR (yellow).

**Supplementary Movie 5**

CryoSparc 3D variability analysis component 0 (top view) showing the GR:Hsp90:FKBP51 focused map density and atomic model. The FKBP51 (purple) proline-rich loop on the FK1 domain displays continuous motion and is dynamically associated with GR (yellow).

**Supplementary Movie 6**

CryoSparc 3D variability analysis component 0 (side view) showing the GR:Hsp90:FKBP51 focused map density and atomic model. The FKBP51 (purple) proline-rich loop on the FK1 domain displays continuous motion and is dynamically associated with GR (yellow).

#### Materials and Methods

*Data analysis and figure preparation*

Figures were created using UCSF Chimera v.1.14[1] and UCSF ChimeraX v1.0.0[2]. GR ligand binding data was analyzed using Prism v.9.4.0 (GraphPad).

*Protein expression and purification*

Human Hsp90α, Hsp70 (gene *Hsp70A1A*), Hop, p23, p23Δhelix (1-112), FKBP51, FKBP52, and yeast Ydj1 (Hsp40) were expressed in the pET151 bacterial expression plasmid with a cleavable N-terminal, 6x-His tag. Human Bag-1 isoform 4 (116-345) was expressed in a pET28a vector with a cleavable N-terminal, 6x-His tag. Proteins were expressed and purified by the following procedure. Proteins were expressed in *E. coli* BL21 star (DE3) strain. Cells were grown in either LB or TB at 37°C until $OD_{600}$ reached 0.6-0.8 and then induced with 0.5 mM IPTG overnight at 16°C. Cells were harvested and lysed in 50 mM Potassium Phosphate pH 8, 500 mM KCl, 10 mM imidazole pH 8, 10% glycerol, 6 mM βME, and Roche cOmplete, mini protease inhibitor cocktail using an EmulsiFlex-C3 (Avestin). Lysate was centrifuged and the soluble fraction was affinity purified by gravity column with Ni-NTA affinity resin (QIAGEN). The protein was eluted with 30 mM Tris pH 8, 50 mM KCl, 250 mM imidazole pH 8, and 6 mM βME. For Hsp90, Hsp70, and Ydj1, an extra wash step with 0.1% Tween20 and 2 mM ATP/$MgCl_2$ was added to the Ni-NTA resin before eluting. The 6x-His tag was removed with TEV protease during the following overnight dialysis in 30 mM Tris pH 8, 50 mM KCl, and 6 mM βME. Cleaved protein was then loaded onto an ion exchange column, MonoQ 10/100 GL (GE Healthcare), with 30 mM Tris pH 8, 50 mM KCl, and 6 mM βME and eluted with a linear gradient of 50-500 mM KCl. Protein was further purified by size exclusion in 30 mM HEPES pH

7.5, 50 mM KCl, 10% glycerol, 1-2 mM DTT using a HiLoad 16/60 Superdex 200 (GE Healthcare) or Hi Load 16/60 Superdex 75 (GE Healthcare). For Hsp70, each peak from ion exchange was collected separately and purified by size exclusion in 30 mM HEPES pH 7.5, 100 mM KCl, 10% glycerol, 4 mM DTT, where only the monomeric peak was then collected. Protein was concentrated, flash frozen, and stored at -80$^{\circ C}$.

*GR DBD-LBD expression and purification*

The human GR DBD-LBD construct contains the GR DNA binding domain (DBD), hinge, and ligand binding domain (LBD) (418-777) with solubilizing mutation F602S. The construct was codon optimized and expressed in the pMAL-c3X derivative with an N-terminal cleavable 6x-His-MBP tag. For datasets I, II, and III, GR DBD-LBD was expressed and purified with ligand as follows. GR DBD-LBD were expressed in *E. coli* BL21 star (DE3) strain. Cells were grown in either LB or TB at 37°C until $OD_{600}$ reached 0.6 and then 100 μM dexamethasone and 50 μM $ZnCl_2$ were added. Cells were induced with 1 mM IPTG at $OD_{600}$ 0.8. Cells were grown overnight (~16-18 hours) at 16°C. Cells were harvested and lysed in 50 mM Tris pH 8, 300 mM KCl, 50 μM dexamethasone, 5 mM imidazole pH 8, 10% glycerol, 2 mM DTT, and 0.2 mM PMSF. Lysate was centrifuged and the soluble fraction was affinity purified by gravity column with Ni-NTA affinity resin (QIAGEN). During Ni-NTA affinity purification, the resin was washed with a buffer containing 30mM Tris pH 8, 500mM KCl, 50 μM dexamethasone, 10% glycerol, 2mM DTT, 2mM ATP, 5mM $MgCl_2$, and 0.1% Tween20. Then the resin was washed with a buffer containing 30mM Tris pH 8, 500mM KCl, 50 μM dexamethasone, 10% glycerol, 2mM DTT, and 5mM EDTA. The protein was eluted with 30 mM Tris pH 8, 150 mM

KCl, 50 μM dexamethasone, 300 mM imidazole pH 8, 10% glycerol, and 3mM DTT. Protein was then purified by size exclusion in 30 mM HEPES pH 7.5, 150 mM KCl, 50 μM dexamethasone, 10% glycerol, and 4 mM DTT using a HiLoad 16/60 Superdex 200 (GE Healthcare). Protein was purified a second time by size exclusion using a HiLoad 16/60 Superdex 200 (GE Healthcare) with the same buffer to further remove degradation products. Protein was concentrated, flash frozen, and stored at -80$^{\circ C}$.

For dataset IV and GR ligand binding assays, apo GR DBD-LBD was expressed and purified as follows. GR DBD-LBD were expressed in *E. coli* BL21 star (DE3) strain. Cells were grown in either LB or TB at 37°C until $OD_{600}$ reached 0.6 and then 180 μM dexamethasone and 50 μM $ZnCl_2$ were added. Cells were induced with 1 mM IPTG at $OD_{600}$ 0.8. Cells were grown overnight (~16-18 hours) at 16°C. Cells were harvested and lysed in 50 mM Tris pH 7.5, 300 mM KCl, 50 μM dexamethasone, 20 mM imidazole pH 8, 10% glycerol, 2 mM DTT, 0.04% CHAPS, 1 mM PMSF, and Roche cOmplete, mini protease inhibitor cocktail using an EmulsiFlex-C3 (Avestin). Lysate was centrifuged and the soluble fraction was affinity purified by gravity column with Ni-NTA affinity resin (QIAGEN). During Ni-NTA affinity purification, the resin was washed with a buffer containing 50mM Tris pH 7.5, 300mM KCl, 50 μM dexamethasone, 20 mM imidazole pH 8, 10% glycerol, 2mM DTT, 2mM ATP, 5mM $MgCl_2$, 0.04% CHAPS, 1 mM PMSF, and Roche cOmplete, mini protease inhibitor cocktail using an EmulsiFlex-C3 (Avestin). The protein was eluted with 30 mM Tris pH 7.5, 150 mM KCl, 100 μM cortisol, 300 mM imidazole pH 8, 10% glycerol, and 3mM DTT. Protein was then dialyzed overnight in a buffer containing 30mM Tris pH 7.5, 150mM KCl, 10% glycerol, and 2mM DTT. Protein was then purified by hydrophobic interaction chromatography (HIC) on a HiScreen Butyl-S FF (4.7mL) column (Cytiva Life Sciences) to remove degradation products. First, solid

KCl was slowly added to the protein solution at 4°C to a final concentration of 2mM KCl. Then the protein was injected onto the HIC column and eluted over a gradient of 2 M- 0 M KCl overs 10 column volumes in a buffer containing 30mM Tris pH 7.5, 10% glycerol, and 2mM DTT. Then the protein was further purified by size exclusion in 30 mM HEPES pH 8, 150 mM KCl, 10% glycerol, and 0.5mM TCEP using a HiLoad 16/60 Superdex 200 (GE Healthcare). Protein was then dialyzed for 3 days, with fresh buffer each day, in a buffer containing 30mM HEPES pH 8, 150 mM KCl, 10% glycerol, and 2mM DTT. Protein was concentrated, flash frozen, and stored at -80$^{\circ C}$.

*GR:Hsp90:FKBP complex sample preparation*

The GR chaperone cycle was reconstituted *in vitro* with purified components as previously described[3]. Buffer conditions were 30 mM HEPES pH 8, 50 mM KCl, 0.05% Tween20, and 2 mM TCEP. Proteins and reagents were added at the following concentration: 5 μM MBP-GR LBD, 2 μM Hsp40, 5 μM Hsp70, 5 μM Hop, 15 μM Hsp90, 15 μM p23, 15 μM FKBP51 or FKBP52, and 5 mM ATP/$MgCl_2$. This reaction was incubated at room temperature for 60 minutes, then 15 μM FKBP51 or FKBP52, 15 μM Bag-1, and 20 mM sodium molybdate (used to stabilize the closed conformation of Hsp90[4-6], likely by acting as a γ-phosphate analog to stabilize the post-ATP hydrolysis transition state of Hsp90 (**Extended Data 3a, 7a)**) were added, and the reaction was incubated at room temperature for another 30 minutes. Following incubation, amylose resin (New England Biolabs) was added to the reactions in a 1:1 ratio and incubated at 4°C with nutation. Resin was then washed 4 times with wash buffer (30 mM HEPES pH 8, 50 mM KCl, 5 mM ATP/$MgCl_2$, 0.05% Tween20, 2 mM TCEP, 20 mM sodium molybdate) and eluted with 50 mM maltose in elution buffer (30 mM HEPES pH 8, 50 mM KCl,

2 mM TCEP, 20 mM sodium molybdate). The elution was analyzed by SDS-PAGE (4-12% acrylamide gel) (**Extended Data Fig. 1c**). The elution was concentrated and purified by size exclusion using a Superdex 200 Increase 3.2/300 (Cytiva Life Sciences) and fractions were analyzed by SDS-PAGE (4-12% acrylamide gel) (**Extended Data 1d**). Fractions containing the full complex were concentrated to ~2 μM and crosslinked with 0.02% glutaraldehyde for 20 minutes at room temperature (**Extended Data Fig. 1c**). 2.5 μL of sample was applied to glow-discharged QUANTIFOIL R1.2/1.3, 400-mesh, copper holey carbon grid (Quantifoil Micro Tools GmbH) and plunge-frozen in liquid ethane using a Vitrobot Mark IV (FEI) with a blotting time of 12-16 seconds, blotting force 3, at 10°C, and with 100% humidity.

*Cryo-EM data acquisition*

All data was acquired using SerialEM software v.4.0[7] and collected on an FEI Titan Krios electron microscope (Thermo Fisher Scientific) operating at 300kV using a K3 direct electron camera (Gatan) and equipped with a Bioquantum energy filter (Gatan) set to a slit width of 20 eV (example micrographs **Extended Data Fig. 1e**). Images were recorded at a nominal magnification of 105,000×, corresponding to a physical pixel size of 0.835 Å. Datasets I, II, and III were collected in super resolution mode, corresponding to a super resolution pixel size of 0.4175 Å. Dataset IV was not collected in super resolution mode. All datasets were acquired using fringe-free imaging (FFI) and multi-hole targeting using image shift in which 3 micrographs were collected per hole. A nominal defocus range of 0.8 μm –2.0 μm under focus was used for all datasets. For exposure, frame rates, and total dose see **Extended Data Table 1**.

Two small datasets on the GR:Hsp90:FKBP52 and GR:Hsp90:FKBP51 complex were collected before the larger datasets described above. The smaller datasets were collected on samples prepared in a similar manner as described above. For GR:Hsp90:FKBP52, images were collected on a FEI Titan Krios electron microscope (Thermo Fisher Scientific) operating at 300kV using a K3 direct electron camera (Gatan). Images were recorded in super-resolution mode at a nominal magnification of 105,000×, corresponding to a physical pixel size of 0.835 Å. A nominal defocus range of 0.8 μm –2.0 μm underfocus was used. A total exposure of 5.9 seconds was used with 0.05 second subframes (117 total frames). A dose rate of 8.0 $e^-$/pix/s was used, with a total dose of 67 $e^-/Å^2$. For GR:Hsp90:FKBP51, images were collected on a Talos Arctica (Thermo Fisher Scientific) operating at 200kV using a K3 direct electron camera (Gatan). Images were recorded in super-resolution mode at a nominal magnification of 28,000×, corresponding to a physical pixel size of 1.44 Å. A nominal defocus range of 0.8 μm –2.0 μm underfocus was used. A total exposure of 11.5 seconds was used with 0.1 second subframes (115 total frames). A dose rate of 10.4 $e^-$/pix/s was used, with a total dose of 57.5 $e^-/Å^2$.

*Cryo-EM data processing*

The smaller GR:Hsp90:FKBP52 dataset consisted of 2,022 dose-fractionated image stacks, which were motion corrected using UCSF MotionCor2[8] and analyzed with RELION v.3.0.8[9]. Motion corrected images were used for contrast transfer function (CTF) estimation using CTFFIND v.4.1[10] and Laplacian-of-Gaussian particle picking was done in RELION. Multiple rounds of 3D classification with symmetry C1 were performed with the GR-maturation complex (PDB ID: 7KRJ)[11] as a low-pass-filtered (20 Å) initial model until a medium-resolution (~8 Å) GR:Hsp90:FKBP52 reconstruction was obtained from 11,756 particles. This reconstruction was used as an initial reference for the larger GR:Hsp90:FKBP52 datasets.

The smaller GR:Hsp90:FKBP51 dataset consisted of 1,181 dose-fractionated image stacks, which were motion corrected using UCSF MotionCor2[8] and analyzed with RELION v.3.0.8[9]. Motion corrected images were used for contrast transfer function (CTF) estimation using CTFFIND v.4.1[10] and Laplacian-of-Gaussian particle picking was done in RELION. Multiple rounds of 3D classification with symmetry C1 were performed with the GR-maturation complex (PDB ID: 7KRJ)[11] as a low-pass-filtered (20 Å) initial model until a medium-resolution (~6 Å) GR:Hsp90:FKBP51 reconstruction was obtained from 45,000 particles. This reconstruction was used as an initial reference for the larger GR:Hsp90:FKBP51 datasets.

Datasets I-IV were motion corrected using UCSF MotionCor2 and analyzed with RELION v.3.1.0. Motion corrected images with dose weighting were used for contrast transfer function (CTF) estimation using CTFFIND v.4.1 and reference-based picking was done in RELION using the corresponding references from the smaller datasets described above. The processing scheme for datasets I-IV are depicted in **Extended Data Fig. 2** and **Extended Data Fig. 6**. After initial rounds of 3D classification with symmetry C1, the GR:Hsp90:FKBP52 datasets (I and II) were combined and GR:Hsp90:FKBP51 datasets (III and IV) were combined.

For the GR:Hsp90:FKBP52 combined dataset, a particle stack of ~496,000 particles was obtained representing a GR:Hsp90:FKBP52 reconstruction at nominal resolution 3.82 Å. This stack was then subjected to 3D classification without alignment and subsequent 3D refinement on the best classes (~307,000 particles), which yielded the best overall consensus reconstruction at a nominal resolution of 3.56 Å. Additionally, to improve the resolution of the GR:FKBP52 region, the ~496,000 particle stack was subjected to signal subtraction of the Hsp90 region. Focused refinement on the Hsp90-subtracted particle stack was then performed (initial angular sampling 1.8°, initial offset range 3 pixels, initial offset step 0.75 pixels, local searches from auto-sampling 1.8°) using a mask including GR and FKBP52 only. Focused classification without alignment was then performed using a mask including GR and FKBP52 only. Focused refinement on the best class (~107,000 particles) was then performed (initial angular sampling 0.9°, initial offset range 3 pixels, initial offset step 0.75 pixels, local searches from auto-sampling 0.9°) using focused refinement using a mask including GR and FKBP52 only, which yielded a GR:FKBP52 reconstruction with a nominal resolution of 4.31 Å.

Per-particle CTF, beam-tilt refinement, trefoil and 4$^{th}$ order aberration refinement, and astigmatism were estimated for both the consensus reconstruction and GR:FKBP52 focused reconstruction in RELION. The corrected particles stacks were then imported to CryoSparc (v3.3.2) and 2D Classification was performed to clean-up the particle stacks. The consensus reconstruction (307,109 particles) was subjected to Non-Uniform Refinement with an envelope mask and a mask including Hsp90 only, each of which refined to a nominal resolution of 3.01 Å and 2.96 Å, respectively. The GR:FKBP52 focused reconstruction (106,318 particles) was subjected to Local Refinement with a mask including GR and FKBP52 only, which refined to a nominal resolution of 3.76 Å.

For the GR:Hsp90:FKBP51 combined dataset, a particle stack of ~500,000 particles was obtained, representing a GR:Hsp90:FKBP51 reconstruction at a nominal resolution of 4.05 Å. This stack was then subjected to 3D classification without alignment and subsequent 3D refinement on the best classes (~172,000 particles) to obtain the best overall consensus reconstruction at a nominal resolution of 4.18 Å. Additionally, to improve the resolution of the GR:FKBP51 region, the ~500,000 particle stack was subjected to signal subtraction of the Hsp90 region. Focused refinement on the Hsp90-subtracted particle stack was then performed (initial angular sampling 0.9°, initial offset range 3 pixels, initial offset step 1 pixels, local searches from auto-sampling 0.9°)  using a mask including GR and FKBP51 only. Focused classification without alignment was then performed using a mask including GR and FKBP51 only. Focused refinement on the best class (~120,000 particles) was then performed (initial angular sampling 0.9°, initial offset range 3 pixels, initial offset step 1 pixels, local searches from auto-sampling 0.9°) using focused refinement using a mask including GR and FKBP51 only, which yielded a GR:FKBP51 reconstruction with a nominal resolution of 4.31 Å.

Per-particle CTF, beam-tilt refinement, trefoil and $4^{th}$ order aberration refinement, and astigmatism were estimated for both the consensus reconstruction and GR:FKBP51 focused reconstruction in RELION. The corrected particles stacks were then imported to CryoSparc (v3.3.2) and 2D Classification was performed to clean-up the particle stacks. The consensus reconstruction (171,778 particles) was subjected to Non-Uniform Refinement with an envelope mask and a mask including Hsp90 only, each of which refined to a nominal resolution of 3.23 Å. The GR:FKBP51 focused reconstruction (109,900 particles) was subjected to Local Refinement with a mask including GR and FKBP51 only, which refined to a nominal resolution of 4.14 Å.

All final reconstructions were post-processed in CryoSparc in which the nominal resolution was determined by the gold standard Fourier shell correlation (FSC) using the 0.143 criterion (**Extended Data Fig. 2, Extended Data Fig. 6**). Maps were sharpened in CryoSparc and filtered to their estimated resolution. A composite map for both GR:Hsp90:FKBP51 and GR:Hsp90:FKBP52 was generated by combining the overall consensus refinement map with the GR:FKBP focused refinement map using vop maximum in Chimera. Note that the composite maps were only used for presentation in **Fig. 1a** and **Fig. 4a**, but not used in atomic model building or refinement.

CryoSparc 3D Variability Analysis was performed for the focused GR:FKBP51 and

GR:FKBP52 reconstructions with the following parameters: number of modes to solve = 3, symmetry = C1, filter resolution = 6 Å, filter order = 1.5, high pass order = 8, per-particle scale = optimal, number of iterations = 20, and lambda = 0.01.

For both the GR:Hsp90:FKBP51 and GR:Hsp90:FKBP52 complexes, no ligand-free GR complexes were identified during image analysis, despite many rounds of focused classification on GR at various stages of data processing. Only classes with clear ligand density in the GR ligand binding pocket were obtained, suggesting ligand-free GR is either too dynamic or quickly released from the complex, consistent with findings during processing of the GR-maturation complex[11].

*Model building and refinement*

For the GR:Hsp90:FKBP52 atomic model, dexamethasone-bound GR LBD and the closed Hsp90 dimer from the GR-maturation complex (PDB ID: 7KRJ)[11] along with the AlphaFold model of human FKBP52 (accession number: AF-Q02790) were used as starting models[12]. Additionally, the Hsp90 MEEVD peptide from the FKBP51:Hsp90 MEEVD crystal structure (PDB ID: 5NJX)[13] was used. For the GR:Hsp90:FKBP51 atomic model, dexamethasone-bound GR LBD and the closed Hsp90 dimer from the GR-maturation complex (PDB ID: 7KRJ)[11] along with human FKBP51 from the Hsp90:FKBP51:p23 cryo-EM structure (PDB ID: 7L7I)[14] were used as starting models. Additionally, the Hsp90 MEEVD peptide from the FKBP51:Hsp90 MEEVD crystal structure (PDB ID: 5NJX)[13] was used.

Models were refined using Rosetta v.3.11 throughout. Following the split map approach[15] to prevent and monitor overfitting, the Rosetta iterative backbone rebuilding procedure was used to refine models against one of the half maps obtained from RELION, with the other half map only used for validations. Structurally uncharacterized regions, including the FKBP52 TPR:Hsp90 CTD interaction, the FKBP51:HSP90 CTD interaction, and the Hsp90 lumen:GR pre-Helix 1 interaction, were built *de novo* into consensus maps or focused maps using RosettaCM[16]. These regions were then further refined using the same Rosetta iterative backbone rebuilding procedure. With a proper density weight obtained using the half maps, the final model of the GR:Hsp90:FKBP52 and GR:Hsp90:FKBP51 complex was refined against the full reconstruction allowing only sidechain and small-scale backbone refinement. The final refinement statistics are provided in **Extended Data Table 1**.

*Fluorescence polarization assays*

Fluorescence polarization of fluorescent dexamethasone (F-dex) (Thermo Fisher) was measured on a CLARIOstar Plus microplate reader (BMG LabTech) with excitation/emission wavelengths of 485/538 nm, and temperature control set at 25°C. Buffer conditions were 50 mM HEPES pH 8, 100 mM KCl, 2 mM DTT. For equilibrium ligand binding in **Fig. 3e,f; 4f,** and **Extended Data Fig. 8a,b**, proteins were pre-equilibrated together at room temperature for 60 minutes prior to F-dex addition. Proteins and reagents were added at the following concentration: 10 nM F-dex, 100 nM GR DBD-LBD, 2 μM Hsp40, 2 μM Bag-1, 15 μM Hsp70, 15 μM Hsp90, 15 μM Hop, 15 μM p23 or $p23_{\Delta helix}$, 15 μM FKBP or FKBP mutants, and 5 mM ATP/$MgCl_2$. Note that the dissociation constant ($K_D$) between GR and F-dex is ~150 nM[3]. Ligand binding was initiated with 10 nM F-dex and association was measured until reaching equilibrium. The plotted equilibrium values in **Fig. 3e,f; 4f** and **Extended Data Fig. 8a,b** represent the mean of 3 biologically independent samples with error bars representing the standard deviation. Polarization values are plotted as the change in polarization from the control sample (10 nM F-dex, 100 nM GR DBD-LBD, 2 μM Hsp40, 2 μM Bag-1, 15 μM Hsp70, 15 μM Hsp90, 15 μM Hop, 15 μM p23, and 5 mM ATP/$MgCl_2$). For equilibrium ligand binding in **Extended Data Fig. 5d**, proteins were pre-equilibrated together at room temperature for 30 minutes prior to F-dex addition. Proteins and reagents were added at the following concentration: 10 nM F-dex, 100 nM GR and 15 μM FKBP51 or FKBP52. Ligand binding was initiated with 10 nM F-dex and association was measured until reaching equilibrium. The plotted data points for each reaction represent 3 biologically independent samples. GR ligand binding behavior was affected by buffer conditions; therefore, reactions were always normalized such that each reaction had equivalent amounts of buffer reagents.

*Sequence alignments*

For the FKBP52 (gene *FKBP4*) sequence alignments in **Fig. 2f**, sequences were obtained from Uniprot[17], aligned in Clustal Omega[18] (https://www.ebi.ac.uk/Tools/msa/clustalo/), and visualized in JalView 2.11.1.0[19]. Sequences in the alignment are: *H. sapiens* FKBP52, *M. musculus* FKBP52, *R. norvegicus* FKBP52, *D. melanogaster* FKBP52, *T. guttata* FKBP52, *G. gallus* FKBP52, *X. tropicalis* FKBP52, and *H. sapiens* FKBP51 (Uniprot accession codes: Q02790, P30416, Q9QVC8, Q6IQ94, H0ZSE5, A0A3Q3B0L8, A0A310SUH5, Q13451 respectively). For **Fig. 2g**, sequences were obtained from Uniprot[17], aligned in Clustal Omega[18] (https://www.ebi.ac.uk/Tools/msa/clustalo/), and conservation scores were calculated and mapped onto GR from the GR:Hsp90:FKBP52 atomic model using UCSF Chimera v.1.14[1]. Sequences in the alignment are: *H. sapiens* GR, *M. musculus* GR, *R. norvegicus* GR, *T. guttata* GR, *G. gallus* GR, *X. tropicalis* GR, *D. rerio* GR (Uniprot accession codes: P04150, P06537, P06536, A0A674H6U9, A0A1D5PRD7, Q28E31, A0A2R8QN75, respectively).

For **Extended Data Fig. 9b**, the sequences were obtained from Uniprot[17], aligned in Clustal Omega[18] (https://www.ebi.ac.uk/Tools/msa/clustalo/), and mapped onto GR from the maturation complex using Chimera v.1.14[1] a. Sequences in the alignment are the human steroid hormone receptors: glucocorticoid receptor, mineralocorticoid receptor, androgen receptor, progesterone receptor, estrogen receptor α and β (Uniprot accession codes: P04150, P08235, P10275, P06401, E3WH19, Q92731, respectively). Conservation was calculated using percent conservation in Chimera v.1.14[1] (with AL2CO[20] parameters (unweighted frequency estimation and entropy-based conservation measurement)).

*Analysis of FKBP52 mutant expression by Western blot*

Wild-type (JJ762) cells expressing empty vector (pRS423GPD), or plasmid-borne wild-type or mutant FKBP52 (pRS423GPD-FKBP52) were lysed and subjected to SDS-PAGE (10% acrylamide gel) followed by immunoblot analysis with a monoclonal antibody specific for FKBP52 (Hi52b, a gift from Dr. Marc Cox, The University of Texas at El Paso)[21] (**Extended Data Fig**. **5a**)  An antibody against PGK1 (Invitrogen #459250) was used as a loading control.

*In vivo GR activity assays*

Relating to **Fig. 2e**, the effect of overexpression of wild-type FKBP52 on GR activity was determined as previously described[21]. GR activity was measured in the wild-type *S. cerevisiae* strain (JJ762) expressing GR on a single copy plasmid (p414GPD-GR) and the GRE-lacZ reporter plasmid pUCDSS-26X.  Wild-type or mutant FKBP52 was expressed in the pRS423GPD plasmid. Cells were grown at 30°C with shaking overnight in selective media, diluted 10-fold and grown to $OD_{600}$ 0.4-0.5.  Cultures were split in two and one set was induced with ligand (50 nM DOC, deoxycorticosterone) (Sigma) for one hour. The β-galactosidase (β-gal) activity of paired samples in the presence and absence of hormone was measured as described using the yeast β-galactosidase assay kit from Thermo Scientific (Catalog number #75768). Assays contained triplicate samples and were conducted at least twice with each mutant. A representative assay is shown.

Fold GR activity was determined by the increase in normalized β-gal activity in the hormone treated sample relative to the untreated paired sample. Relative GR activation was calculated by normalizing the fold GR activity of each sample to the average fold GR activity of strain JJ762 expressing p423GPD (empty vector [e.v.]). The fold increase in GR activities compared to the empty vector (e.v.) control are shown (mean±SD).

*Quantification and statistical analysis*

All data were tested for statistical significance with Prism v.9.4.0 (GraphPad) (n.s. $P \geq 0.05$; * $P \leq 0.05$; ** $P \leq 0.01$; *** $P \leq 0.001$; **** $P \leq 0.0001$). Statistical details (including sample sizes (*n*), F-statistics, p-values, and degrees of freedom) are included in the figure legends for each experiment where possible. Relating to **Fig. 2e**, significance was evaluated using a one-way ANOVA ($F_{(6,14)} = 67.82$; $p < 0.0001$) with *post-hoc* Dunnett's multiple comparisons test P-values: p(e.v. vs. 52) < 0.0001, p(52 vs. 52ΔFK1) < 0.0001, p(52 vs. 52 S118A) < 0.0001, p(52 vs. 52 Y161D) = 0.0001. Relating to **Fig 3e**, significance was evaluated using a one-way ANOVA ($F_{(3,8)} = 541.2$; $p < 0.0001$) with *post-hoc* Šídák's test. P-values: p(Chaperones vs. Chaperones + 52) = 0.0002, p(Chaperones + 52 vs. Chaperones w/ p23Δhelix + 52) < 0.0001, p(Chaperones w/ p23Δhelix vs. Chaperones w/ p23Δhelix + 52) < 0.0001. Relating to **Fig. 3f,** significance was evaluated using a one-way ANOVA ($F_{(5,12)} = 761.5$; $p < 0.0001$) with *post-hoc*

Šídák's test. P-values < 0.0001 for each comparison. Relating to **Fig. 4f** statistical significance was evaluated by an ordinary one-way ANOVA with *post-hoc* Šídák's multiple comparisons test. P-values: p(Chaperones vs. Chaperones + w/ p23Δhelix) < 0.0001, p(Chaperones vs. Chaperones w/ p23Δhelix + 51) = 0.0287, p(Chaperones w/ p23Δhelix + 51 vs. Chaperones w/ p23Δhelix + 51 L119P) < 0.0001, p(Chaperones w/ p23Δhelix + 51 vs. Chaperones w/ p23Δhelix + 52) < 0.0001, p(Chaperones w/ p23Δhelix + 52 vs. Chaperones w/ p23Δhelix + 52 P119L) < 0.0001. Relating to Extended Data Fig. 5d, statistical significance was evaluated by an ordinary one-way ANOVA ($F_{(2,6)}$ = 1414, p < 0.0001) with *post-hoc* Tukey's multiple comparisons test. P-values: p(GR vs. GR + 51) < 0.0001, p(GR vs. GR + 52) < 0.0001, p(GR + 51 vs. GR + 52) = 0.004. Relating to **Extended Data Fig. 5d**, statistical significance was evaluated by an ordinary one-way ANOVA *post-hoc* Tukey's multiple comparisons test using Prism v.9.4.0 (GraphPad).Relating to **Extended Data Fig. 8a** statistical significance was evaluated by an unpaired two-tailed t test, p-value =0.5737. Relating to **Extended Data Fig. 8b**, statistical significance was evaluated by an ordinary one-way ANOVA ($F_{(6,12)}$ = 647.1, p < 0.0001) with *post-hoc* Šídák's multiple comparisons test. P-values: p(Chaperones vs. Chaperones -p23) < 0.0001, p(Chaperones vs. Chaperones -p23 + 51) < 0.0001= 0.0287, p(Chaperones -p23 vs. Chaperones -p23 + 51) = 0.0123, p(Chaperones + Mo. Vs. Chaperones -p23 + Mo. ) < 0.0001, p(Chaperones + Mo. Vs. Chaperones -p23 + 51 + Mo. ) = 0.1640, p(Chaperones -p23 + Mo. Vs. Chaperones -p23 + 51 + Mo. ) < 0.0001.

**Data availability**

The cryo-EM maps generated in this study have been deposited in the Electron Microscopy Data Bank (EMDB) under the accession codes EMD-29068 (GR:Hsp90:FKBP52) and EMD-29069

(GR:Hsp90:FKBP51). The atomic coordinates have been deposited in the PDB under the accession code 8FFV (GR:Hsp90:FKBP52) and 8FFW (GR:Hsp90:FKBP51). Publicly available PDB entries used in this study are: 7KRJ, 5NJX, 7L7I, 1M2Z, 4LAV, 6TXX and AlphaFold AF-Q02790. Protein sequence data for sequence alignments are available from Uniprot (see **Methods** for accession codes).

#### Methods References


1 Pettersen, E. F. *et al.* UCSF Chimera--a visualization system for exploratory research and analysis. *J Comput Chem* **25**, 1605-1612, doi:10.1002/jcc.20084 (2004).

2 Goddard, T. D. *et al.* UCSF ChimeraX: Meeting modern challenges in visualization and analysis. *Protein Sci* **27**, 14-25, doi:10.1002/pro.3235 (2018).

3 Kirschke, E., Goswami, D., Southworth, D., Griffin, P. & Agard, D. Glucocorticoid Receptor Function Regulated by Coordinated Action of the Hsp90 and Hsp70 Chaperone Cycles. *Cell* **157**, 1685-1697, doi:10.1016/j.cell.2014.04.038 (2014).

4 Johnson, J. L. & Toft, D. O. Binding of p23 and hsp90 during assembly with the progesterone receptor. *Mol Endocrinol* **9**, 670-678, doi:10.1210/mend.9.6.8592513 (1995).

5 Verba, K. A. *et al.* Atomic structure of Hsp90-Cdc37-Cdk4 reveals that Hsp90 traps and stabilizes an unfolded kinase. *Science* **352**, 1542-1547, doi:10.1126/science.aaf5023 (2016).

6 Csermely, P. *et al.* Atp Induces a Conformational Change of the 90-Kda Heat-Shock Protein (Hsp90). *Journal of Biological Chemistry* **268**, 1901-1907 (1993).

7 Schorb, M., Haberbosch, I., Hagen, W. J. H., Schwab, Y. & Mastronarde, D. N. Software tools for automated transmission electron microscopy. *Nat Methods* **16**, 471-477, doi:10.1038/s41592-019-0396-9 (2019).

8 Zheng, S. Q. *et al.* MotionCor2: anisotropic correction of beam-induced motion for improved cryo-electron microscopy. *Nat Methods* **14**, 331-332, doi:10.1038/nmeth.4193 (2017).

9 Scheres, S. H. RELION: implementation of a Bayesian approach to cryo-EM structure determination. *J Struct Biol* **180**, 519-530, doi:10.1016/j.jsb.2012.09.006 (2012).

10 Rohou, A. & Grigorieff, N. CTFFIND4: Fast and accurate defocus estimation from electron micrographs. *J Struct Biol* **192**, 216-221, doi:10.1016/j.jsb.2015.08.008 (2015).

11 Noddings, C. M., Wang, R. Y., Johnson, J. L. & Agard, D. A. Structure of Hsp90-p23-GR reveals the Hsp90 client-remodelling mechanism. *Nature* **601**, 465-469, doi:10.1038/s41586-021-04236-1 (2022).

12 Jumper, J. *et al.* Highly accurate protein structure prediction with AlphaFold. *Nature*, doi:10.1038/s41586-021-03819-2 (2021).

13 Kumar, R., Moche, M., Winblad, B. & Pavlov, P. F. Combined x-ray crystallography and computational modeling approach to investigate the Hsp90 C-terminal peptide binding to FKBP51. *Sci Rep* **7**, 14288, doi:10.1038/s41598-017-14731-z (2017).

14 Lee, K. *et al.* The structure of an Hsp90-immunophilin complex reveals cochaperone recognition of the client maturation state. *Mol Cell* **81**, 3496-3508 e3495, doi:10.1016/j.molcel.2021.07.023 (2021).

15 Wang, R. Y. *et al.* Automated structure refinement of macromolecular assemblies from cryo-EM maps using Rosetta. *Elife* **5**, doi:10.7554/eLife.17219 (2016).

16 Song, Y. *et al.* High-resolution comparative modeling with RosettaCM. *Structure* **21**, 1735-1742, doi:10.1016/j.str.2013.08.005 (2013).

17 UniProt, C. UniProt: the universal protein knowledgebase in 2021. *Nucleic Acids Res* **49**, D480-D489, doi:10.1093/nar/gkaa1100 (2021).

18 Madeira, F. *et al.* The EMBL-EBI search and sequence analysis tools APIs in 2019. *Nucleic Acids Res* **47**, W636-W641, doi:10.1093/nar/gkz268 (2019).

19 Waterhouse, A. M., Procter, J. B., Martin, D. M., Clamp, M. & Barton, G. J. Jalview Version 2--a multiple sequence alignment editor and analysis workbench. *Bioinformatics* **25**, 1189-1191, doi:10.1093/bioinformatics/btp033 (2009).

20 Pei, J. & Grishin, N. V. AL2CO: calculation of positional conservation in a protein sequence alignment. *Bioinformatics* **17**, 700-712, doi:10.1093/bioinformatics/17.8.700 (2001).

21 Riggs, D. L. *et al.* The Hsp90-binding peptidylprolyl isomerase FKBP52 potentiates glucocorticoid signaling in vivo. *The EMBO Journal* **22**, 1158-1167, doi:10.1093/emboj/cdg108 (2003).
